# Type I IFN reprograms CD4⁺ T cell lipid metabolism as an antiviral effector mechanism during acute HIV/SIV infection

**DOI:** 10.64898/2026.02.12.705596

**Authors:** Jinhee Kim, Anoop Ambikan, Jakob Harrison-Gleason, Kayla L. Yerlioglu, Iva Filipovic, Romaila Abdelraouf, Catarina Ananias-Saez, Erin B Taylor, Mariluz Arainga, Deepanwita Bose, Francois J Villinger, Ujjwal Neogi, Elena Martinelli

## Abstract

Cellular metabolism regulates HIV/SIV replication and reservoir establishment, yet how infection and antiretroviral therapy initiation (ARTi) shape CD4⁺ T cell metabolism *in vivo* remains poorly defined. Using the SIVmac239 macaque model, we integrated single-cell metabolic profiling (MIST), transcriptomics, lipidomics and genome-scale metabolic modeling to characterize their metabolic remodeling.

At peak viremia, CD4⁺ T cells exhibited shutdown of de novo fatty-acid (FA) synthesis, reflected by acetyl-CoA carboxylase-1 (ACC1) downregulation, inhibition of lipid-anabolic reactions, and depletion of membrane phospholipids. This state was driven by type I interferon (IFN-I) responses, and IFN-I suppressed ACC1 *in vitro*. Pharmacologic inhibition of FA synthesis enhanced T cell activation and exerted direct antiviral effects. Following ARTi, most metabolic pathways in CD4^+^ T cells were suppressed, whereas oxidative phosphorylation (OXPHOS) remained elevated and its levels in effector memory cells correlated with cell-associated-vDNA These findings identify IFN-driven FA synthesis suppression as a novel effector mechanism during acute viral infection, and persistent OXPHOS in effector CD4⁺ T cells as a metabolic correlate of early reservoir establishment.

## Introduction

Cellular metabolism is a critical determinant of Human Immunodeficiency Virus (HIV) and its simian counterpart (SIV) replication and persistence^1, 2^. Over the past decade, mounting evidence has revealed cellular metabolism as a defining feature of infection susceptibility, replication efficiency, and reservoir establishment^3, 4, 5^. HIV preferentially targets metabolically active CD4^+^ T cells exhibiting enhanced glycolysis and oxidative phosphorylation (OXPHOS), independent of activation marker expression^5^. *In vitro* HIV infection triggers coordinated upregulation of multiple metabolic pathways, including enhanced glycolysis, increased OXPHOS, and altered lipid and amino acid metabolism^3, 6^. Glutaminolysis has emerged as particularly crucial to HIV replication, which also requires deoxynucleoside triphosphates for reverse transcription and lipids for membrane biogenesis^3, 6^. However, how HIV infection shapes cellular metabolism of both productively infected cells and bystander CD4^+^ T cells *in vivo* remains largely uncharacterized.

The period from acute infection to antiretroviral therapy (ART) initiation represents a critical window that shapes viral replication dynamics and the establishment of long-lived infected cell populations ^7, 8, 9^. Although the HIV reservoir starts seeding within few days after infection^10, 11^ and continues until viral replication is blocked by ART^12, 13^, the infected CD4⁺ T cell pool undergoes rapid turnover during untreated infection^8, 14^. Hence, several studies suggest that many infected cells that persist on ART are seeded near the time of therapy initiation^7, 8, 9^, highlighting the importance of understanding the changes in cellular metabolism during this transition. Despite its importance, CD4⁺ T cell metabolism during acute infection and ART initiation remains poorly characterized *in vivo*.

Lipid metabolism is known to influence both viral replication and antiviral immunity^15^. Early *in vitro* studies suggested that HIV upregulates fatty acid synthase (FASN)^16^, leading to the assumption that lipid synthesis increases during acute infection. However, more recent work suggested that type I interferons (IFN-I) may reduce fatty acid synthesis in some contexts during viral infections ^17, 18^. Whether HIV/SIV infection in vivo alters FA synthesis or broader cellular lipid metabolism in CD4^+^ T cells, and how such changes influence viral replication dynamics, remains unknown. Notably, ACC1-mediated FA synthesis has been primarily studied in the context of T cell fate decisions, where its inhibition promotes memory formation and regulatory T cell differentiation^19,20^. Whether ACC1 and FA synthesis are modulated during acute viral infection in vivo, what signals drive such modulation, and what consequences this carries for antiviral effector responses, have not been investigated.

Finally, while one study linked cellular OXPHOS activity with viral set point during acute infection^21^, the impact of treated infection with its decreasing antigenic burden on metabolic pathways within T cells and their functional implications has not been investigated.

The present study addresses these gaps by leveraging the SIVmac239 non-human pr^20^imate (NHP) model and a high-dimensional flow cytometry-based approach (MIST - Metabolic Immune Signature Technique) to comprehensively profile metabolic changes in CD4^+^ T cells during acute infection and early ART. Moreover, by integrating single-cell metabolic enzyme profiling with transcriptomics and plasma metabolomics, this work generated the first genome-scale metabolic model (GEM) of SIV infection in macaques. GEMs provide a mechanistic, genome-resolved framework for predicting metabolic states and identifying targets to reverse disease-associated metabolic perturbations.^22, 23^.

We found that acute SIV induces a marked shutdown of lipid anabolic pathways, particularly de novo FA synthesis, coinciding with strong IFN-I responses and resulting in broad depletion of membrane lipids. ART initiation (ARTi) partially restored lipid metabolism, whereas most other metabolic pathways became progressively downregulated during chronic infection on ART. OXPHOS emerged as a notable exception, remaining elevated compared with pre-infection levels. Together, these findings reveal distinct metabolic states across acute infection and ART initiation and identify IFN-driven FA synthesis suppression during acute infection and persistent OXPHOS after treatment initiation as key features of CD4⁺ T cell metabolism in HIV/SIV infection.

## Results

### Peak SIV Infection Leads to Fatty Acids Synthesis Shutdown in CD4^+^ T cells

To understand the impact of SIV infection and ART initiation on CD4^+^ T cells metabolism, we examined PBMC one week before SIVmac239M2 intravenous infection of 8 Indian rhesus macaques (Macaca Mulatta; females) and at weeks 2, 6, 15, and 23 post-infection (pi) (Figure 1A). Week 2 pi represented the time of peak plasma viral load (pVL; median 3.6×10^7^ copies/ml; 95%CI 2×10^8^ – 1.9×10^7^; Figure 1B). All macaques started a fully suppressive ART regimen, including Tenofovir, Dolutegravir, and Emtricitabine at week 3 pi. By week 6 pi, the viral load was already suppressed (median 650 copies/ml; 95%CI 1500 – 25; Fig. 1B), and by week 23 pi pVL was below detection limits (15 copies/ml) in all animals, except A19X093 with 25 copies/ml. At weeks 2, 6, 15 and 23 pi, we also measured intact and total provirus and cell-associated SIV-RNA in PBMC (gag and tat/rev; Figure 1B and Figure S1). PBMC metabolism was first probed by measuring the levels of several rate-limiting enzymes and markers of major metabolic reactions in immune cells by high-dimensional spectra flow cytometry in an assay that we named MIST (<u>m</u>etabolic immune <u>s</u>ignature technique). This approach was similar to the one described as Met-Flow in Ahl PJ et al^24^. The panel (Table S1) was validated on isolated naïve macaque CD4^+^ T cells and PBMC, activated or not with anti-CD3 and anti-CD28 antibodies for 3 days. The assay recapitulated previously reported findings^24^ and the expected increases in ACC1^25, 26^ (Acetyl-CoA Carboxylase 1, the rate limiting enzyme for fatty acid synthesis), CPT1^26^ (Carnitine palmitoyltransferase 1, fatty acid metabolism), HK1 (hexokinase 1, glycolysis)^27^, PRDX2 (Peroxiredoxin-2, oxidative stress)^28^, phospho-RPS6 (Ribosomal protein S6, downstream of mTOR)^29^ as well as classical activation markers including CD25 and CD95^30^ (Figure S2A). In contrast, CD62L (L-selectin) was reduced^31^. Of note, within T cell subsets, correlation with metabolic markers recapitulated expected patterns, including an inverse correlation between naïve CD4^+^ T cells and most metabolic markers upregulated during T cell activation (Figure S2B; gating strategy in Figure S2C). In contrast, the levels of ATP5A and TCF1 expression (ATP5F1A, mitochondrial ATP synthase levels as marker of OXPHOS activity and transcription factor T-cell factor 1, marking quiescent stem-like cells) positively correlated with naive CD4^+^ T cells, while effector memory and TEMRA (terminally differentiated) T cells positively correlated with markers of oxidative stress (PRDX2) and fatty acid oxidation (CPT1A) (Figure S2B). Similar but more granular results were obtained when each CD4^+^ T cell subset was analyzed separately (Figure S2D). Finally, metabolic enzymes and markers upregulated by anti-CD3/CD28 stimulation correlated positively with T cell activation markers and negatively with markers of naïve or quiescent T cells (Figure S2E).

**Fig. 1.**
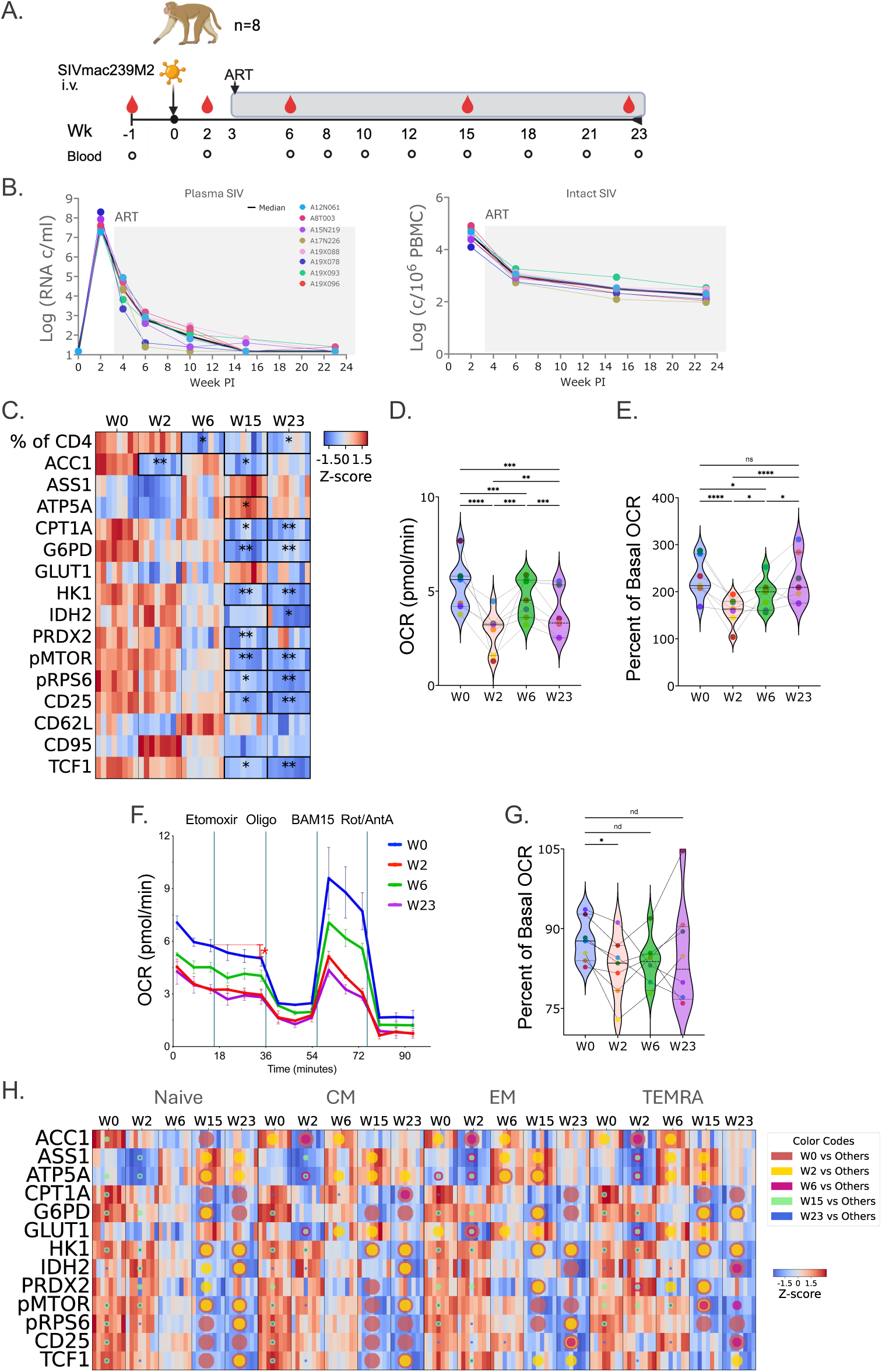
SIV infection and ART lead to metabolic reprogramming of CD4 T cells in vivo. A) Schematic of the NHP study. Eight macaques were infected i.v. with SIVmac239M2. Blood was collected before infection, and weekly or every other week. Suppressive cART started at week 3p.i. PBMC and plasma were analyzed at weeks -1, 2, 6, 15 and 23pi as indicated (red drop). B) Plasma viral loads (left) and levels of intact SIV DNA in PBMC (right) are shown. C) Heatmap of MIST variables’ levels in CD4^+^ T cells at indicated weeks before and after infection (W0= pre-infection bleed) (*q≤0.05; **q≤0.01; Friedman Test with Nemenyi posthoc and TSBH-FDR correction comparing each time point to before infection) D-G) Mitochondrial respiration of sorted CD4⁺ T cells analyzed using a modified Seahorse T Cell Fitness Assay performed in low-glucose to promote fatty-acid (FA) utilization. D) Basal oxygen consumption rate (OCR) is shown in CD4⁺ T cells at different time point post-infection. (*q≤0.05 mixed-effect model with BH-FDR correction) E) Spare respiratory capacity (SRC; maximal OCR − basal OCR) is shown as percent of OCR (*p ≤0.05). F) Representative OCR kinetics from a single animal showing sequential injection of etomoxir, oligomycin, BAM15, and rotenone/antimycin A. G) Etomoxir response (OCR after etomoxir – basal OCR) shown as percent of basal OCR (red asterisk in panel F) (*q≤0.05; Friedman test with BH-FDR correction). H) Heatmap of MIST marker levels in CD4⁺ T cell subsets at the indicated weeks. For visualization, gMFI values for each marker were z-score normalized independently within each subset across all animals and time points, so color intensity reflects longitudinal changes within each subset. Statistical comparisons were performed within each subset using the Friedman test with Nemenyi post-hoc and TSBH-FDR correction comparing each time point to all others. Significant marker–time-point comparisons (q ≤ 0.05) are overlaid as colored circles, where circle color and size denote the comparator time point (W0, W2, W6, W15, or W23).

MIST analysis of CD4^+^ T cells collected before and after SIV infection led to two major observations: a profound decrease in ACC1 levels at peak infection and a broad disruption of major metabolic pathways during prolonged ART compared to pre-ART levels (Figure 1C and representative flow plots in Figure S3). Notably, ACC1, which catalyzes the first committed step in the *de novo* FA synthesis pathway, was markedly reduced at peak viral replication, suggesting an infection-driven metabolic reprogramming of CD4^+^ T cells at the level of lipid metabolism that would lead to decreased FA synthesis. This finding was confirmed by RT-dPCR in a subset of macaques and Western blot in an independent macaque cohort infected with the same viral stock (Figure S4). ACC1 levels partially recovered following ART initiation (W6 pi; Figure 1C); however, they decreased again with prolonged ART (W15 pi).

To functionally confirm that the ACC1 downregulation at peak viral replication resulted in a decrease in FA synthesis and a depletion of FA in CD4^+^ T cells, we ran a modified version of the Seahorse Agilent T cell fitness assay on CD4^+^ T cells isolated from 5 of the 8 animals at W0, W2, W6 and W23 pi. The assay was run in media with limited glucose to stimulate endogenous FA usage and in the presence of etomoxir, which inhibits FA oxidation. Although basal respiration was reduced at all post-infection time points (Figure 1D) and spare respiratory capacity decreased at peak infection before recovering with late ART (Figure 1E), the etomoxir response was calculated separately as the acute change in OCR immediately before versus after etomoxir injection, normalized to basal OCR. This normalized etomoxir response was significantly reduced at W2 (Figure 1F–G), indicating decreased FA-dependent respiration at peak infection. Together, these data support reduced endogenous FA availability and FA utilization in CD4⁺ T cells, consistent with ACC1 downregulation by MIST.

### Treated SIV Infection Leads to Shutdown of Major Metabolic Pathways with the exception of OXPHOS

The levels of several other key metabolic enzymes, including CPT1A (fatty acid oxidation), G6PD (Glucose-6-phosphate dehydrogenase, the rate limiting enzyme in the pentose phosphate pathway, PPP), HK1 (hexokinase 1, the first rate limiting enzyme in glycolysis), PRDX2 (Peroxiredoxin 2, a key enzyme that maintains cellular redox homeostasis), and metabolic regulators, including phospho-MTOR (activated form of the mammalian target of rapamycin, phosphorylated at ser2448 and the downstream pRPS6 (phosphoS6 ribosomal protein phosphorylated at Ser 240/244) significantly decreased in CD4^+^ T cells in late ART (W15 and W23pi) compared to pre-ART levels (Figure 1C). In contrast, ATP5A levels were elevated at W15pi compared to pre-infection levels and did not decrease following ART initiation (Figure 1C). Together, these findings suggest that chronic, ART treated SIV infection leads to a broad suppression of several major metabolic pathways, while mitochondrial respiration (OXPHOS) is preserved or upregulated with ART.

To demonstrate the relevance to human and HIV-1 infection of our NHP findings, we performed a reanalysis of published PBMCs’ single-cell RNA sequencing (scRNA-Seq) data from untreated and ART-treated people with HIV (PWH) compared to PBMC from PWoH (PRJNA662927 and PRJNA681021)^32, 33^. We performed single cell genome-scale metabolic modelling (GEM) and single cell flux balance analysis of naive and central memory CD4^+^ T cells. In line with the NHP data, in treated HIV-1 infection, we found an increased positive flux in complex IV and V compared to untreated and PWoH (Figure S5A). Moreover, we found higher expression of ATP5A in naive CD4^+^ T cells from treated compared to untreated PWH (Figure S5B). Together, the macaque SIV model and human HIV-1 single-cell meta-analysis demonstrated that chronic, ART-treated infection selectively preserves mitochondrial OXPHOS activity, reflected by sustained Complex IV/V flux and ATP5A expression across species.

Of note, comparisons of MIST data across all time points revealed an increase in argininosuccinate synthase 1 (ASS1)—the rate-limiting enzyme for de novo arginine synthesis—during ART (W15 pi) compared with peak infection (Figure S6). In parallel, we observed marked reductions in activated p-mTOR, p-RPS6, CD25, and TCF1 during late ART (week 23 pi) relative to both pre-infection and peak infection levels (Figure S6A).

Subset-specific analyses confirmed and refined these findings (Figure 1H; gating strategy in Figure S7A). Subset frequencies remained largely stable across infection and ART (Figure S7B), indicating that bulk metabolic changes reflected similar within-subset cell-intrinsic remodeling rather than shifts in subset composition. ACC1 downregulation at W2 was significant in memory subsets (CM, EM and TEMRA), but not in naïve CD4⁺ T cells (Figure 1H). The significance of this decrease in the bulk analysis despite a tendency of naïve cells to increase in W2 (Figure S7B) further supports ACC1 decrease being a coordinate metabolic shift in all CD4^+^ T cell subsets. The ATP5A increase between W2 and W6 was significant in memory subsets, but not in naïve cells. In this case, despite the increased proportion of CM at W6 compared to W2, the significance was lost in the bulk analysis. However, by W15 ATP5A was elevated across all subsets, and in the bulk analysis (Figure S6A), indicating that persistent OXPHOS becomes a generalized feature with prolonged ART. ASS1 followed a similar pattern with a significant increase in terminally differentiated subsets EM and TEMRA at W6 compared to W2, which became significant in all subsets and bulk analysis at W15 compared to W2 pi. Finally, ART/decreased viral replication-associated suppression of other major metabolic pathways, including downregulation of HK1, G6PD, PRDX2, pMTOR and pRPS6, was present across all subsets at W15 and W23 compared with pre-infection and peak viral replication levels.

In CD4⁻ T cells, ACC1 expression did not significantly decline at peak viral replication, as observed in CD4⁺ T cells. However, prolonged ART induced a broad metabolic shutdown in CD4^-^T cells, with relative preservation of OXPHOS, albeit to a lesser extent than in CD4⁺ T cells (Figure S6B-C).

### Genome Scale Metabolic modeling and Lipidomics Confirm SIV- and ART-driven Metabolic Reprogramming of CD4^+^ T cells

To independently validate the findings above and gain additional insight into CD4⁺ T cell metabolism, we performed plasma metabolomics and RNA-seq analysis of sorted CD4⁺ T cells at W0, W2, W6, and W23 pi. Unsupervised time-series clustering analysis of the metabolomics data identified two main clusters: one showing a modest increase and the other a decrease at W23 relative to pre-infection levels (Figure 2A–B). The decreased cluster was enriched for metabolites involved in amino-acid metabolism and contained a smaller proportion of lipid metabolites compared with the increased cluster (Figure S8).

**Fig. 2.**
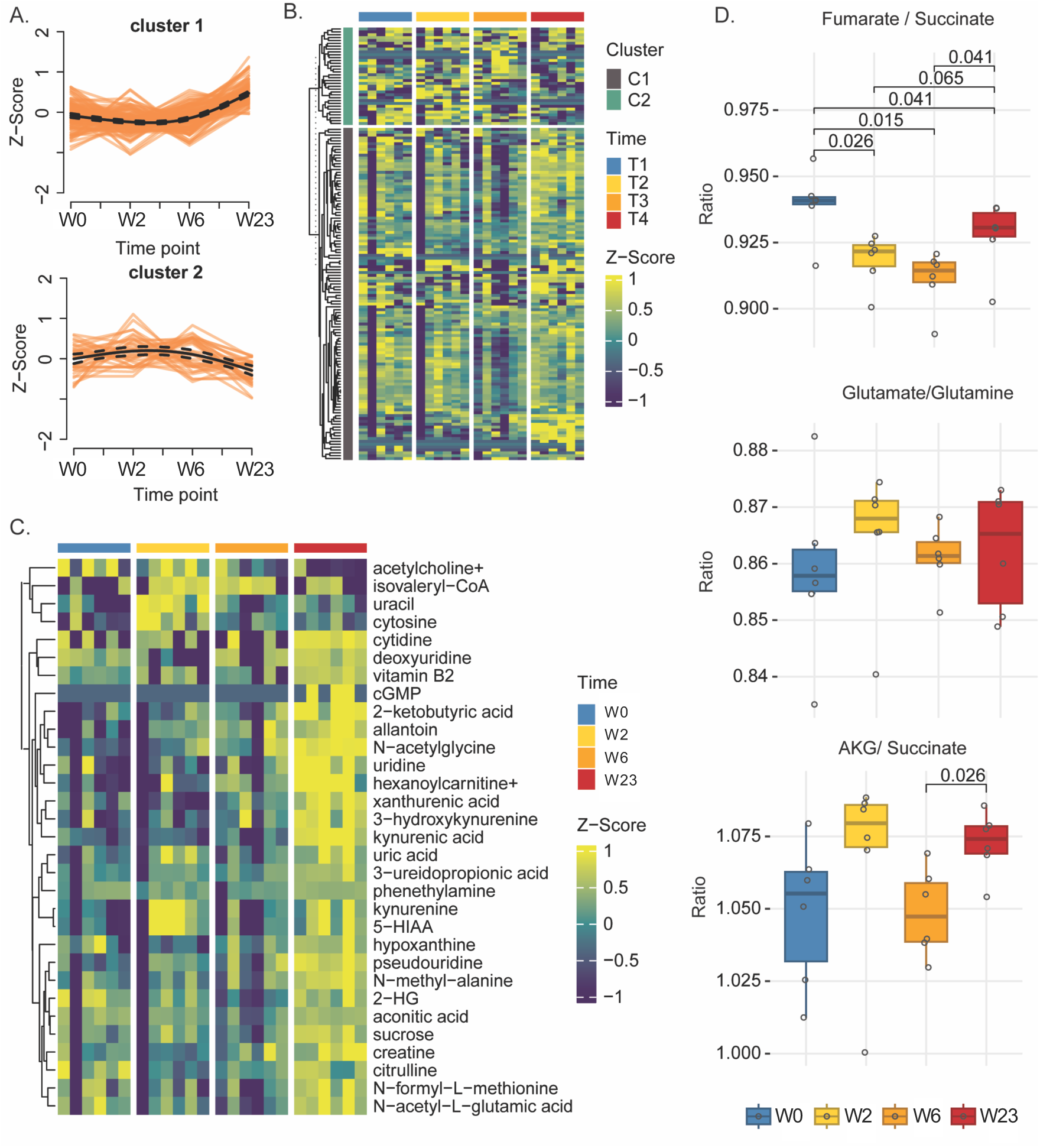
Plasma Metabolomics Suggests Disruption of TCA Cycle with SIV infection. A-B) Tmixclust analysis of plasma metabolomics data at T1 (pre-infection, week -1), T2 peak infection (week 2 pi), T3 (week 6pi, week 3 post-ART initiation) and T4 (week 23pi, week 20 post-ART initiation). Two clusters were identified their relative change is shown as average (A) and as metabolite heatmap per subject (B). C) Time series analysis (Friedmann Test) identified as changing with nominal p<0.05 are shown. D) Ratio of TCA cycle metabolites are shown and compared at different time point by Mann-Whitney test (* p<0.05)

Time-series analysis identified 31 metabolites that were significantly modulated across the four time points (p≤0.05; adj p≤0.2; Figure 2C). Among these, acetylcholine levels declined during late ART, whereas multiple kynurenine pathway metabolites—including kynurenine, kynurenic acid, 3-hydroxykynurenine, and xanthurenic acid—increased, consistent with a shift of tryptophan metabolism toward the kynurenine pathway. In addition, analysis of TCA-cycle metabolites as ratios revealed altered mitochondrial function at peak infection, marked by a sustained reduction in the fumarate/succinate ratio that only partially recovered with ART, indicating persistent dysregulation of the TCA cycle (Figure 2D).

Unsupervised time-series clustering analysis of CD4⁺ T cell RNA-seq data identified three distinct expression clusters. Cluster 1 increased at peak infection and was enriched for interferon (IFN)-stimulated and antiviral response genes, whereas Cluster 2 comprised metabolic genes that were markedly downregulated at peak infection (W2pi), partially recovered with early ART, and declined again during prolonged ART (Figure 3A–C).

**Fig. 3.**
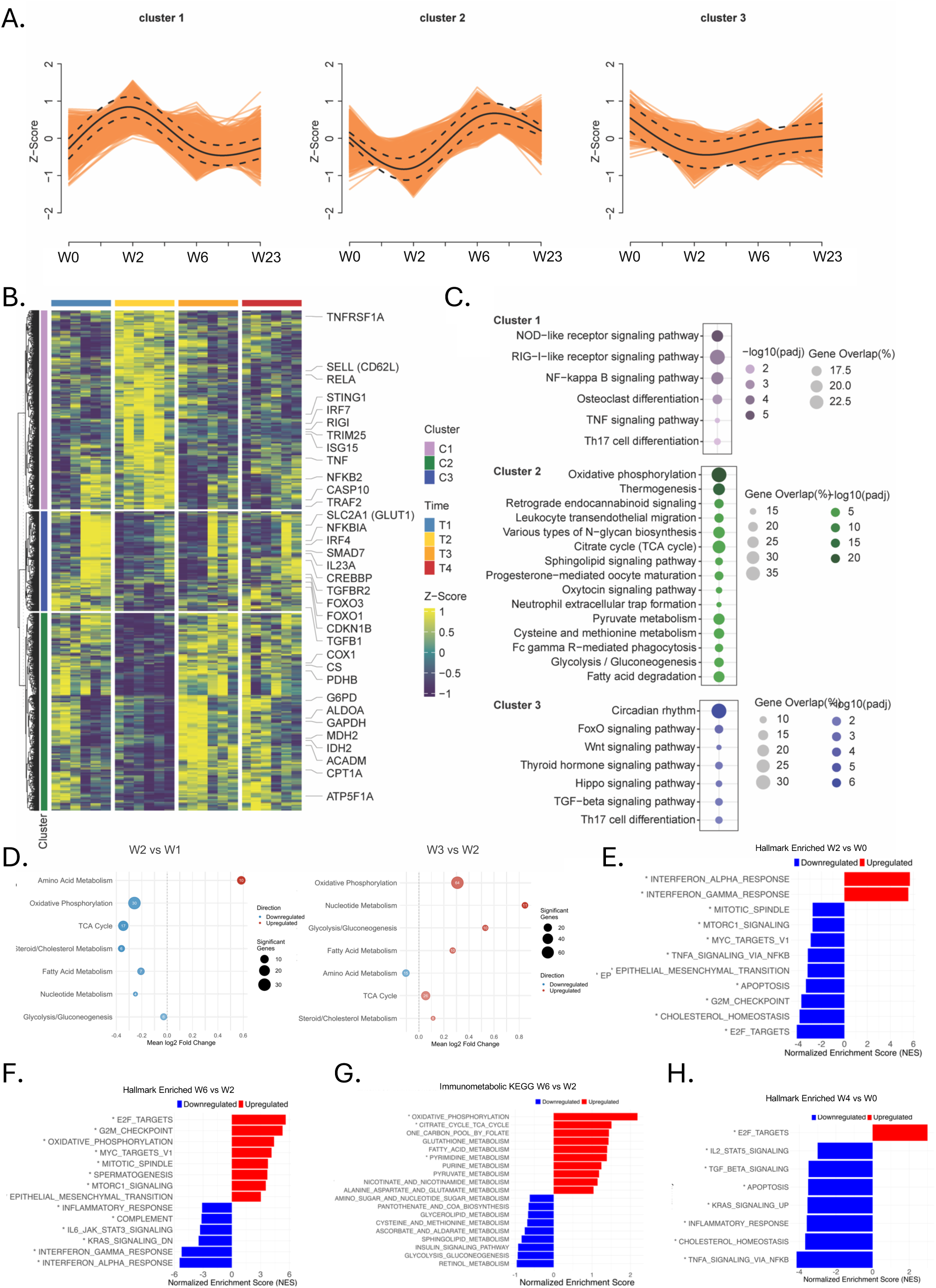
Transcriptomics Data Confirm Antiviral Responses with Decrease Cholesterol metabolism at Peak Infection. A-C) TMixclust analysis of the CD4^+^ T cells RNAseq data revealed 3 clusters. Relative changes in each cluster at the different time post infection (T1 pre-infection, T2 peak infection, T3 week 3 post-ART initiationand T4 week 20 post-ART initiation) are shown (A). Heatmaps of gene-expression changes in each cluster at all time points (B) and pathway-enrichment analyses for each cluster (C) are shown. D-H) DESeq2 differential-expression analyses and pathway-enrichment results for CD4⁺ T cells. D) Bubble plots summarizing metabolic pathway changes in the T2 vs T1 and T3 vs T2 comparisons. Each bubble represents a metabolic pathway, plotted by the mean log₂FC of DEGs (Adj p≤0.05) within that pathway; bubble size denotes the number of DEGs in that comparison, and bubble color reflects the net direction of change (blue, downregulated; red, upregulated). E-F) GSEA performed on genes ranked by log₂FC from the T2 vs T1 and T3 vs T2 comparisons using the Hallmark gene sets (*Adj p≤0.05). G) GSEA performed on genes ranked by log₂FC from the T3 vs T2 comparison using the Immunometabolic KEGG gene sets (*Adj p≤0.05)

Consistent with these patterns, differential gene expression analysis (BH-FDR adj *p*≤0.05) comparing W2 to pre-infection (W0) revealed widespread suppression of genes involved in major energy-producing pathways at peak infection, with the exception of amino acid metabolism (Figure 3D). Most of these pathways showed partial recovery following ART initiation (Figure 3D). Gene set enrichment analysis (GSEA) using the Hallmark collection further supported these findings, identifying IFN responses as the most upregulated pathways and cholesterol homeostasis among the most downregulated at peak infection relative to pre-infection (Figure 3E). To refine these observations, we performed a metabolically focused GSEA using curated KEGG metabolic pathways relevant to immune cells (basic and extended immunometabolic KEGG gene sets; Tables S2 and S3). This analysis identified biosynthesis of unsaturated fatty acids as the most downregulated pathway at peak SIV infection, consistent with the MIST and Seahorse data (Figure S9A). In contrast, Hallmark GSEA comparing early ART (W6) to peak infection (W2) demonstrated a broad increase in metabolic activity, including enrichment of G2M checkpoint, OXPHOS, MYC targets (V1), and MTORC1 signaling pathways, alongside a concurrent reduction in IFN-related pathways (Figure 3F). Metabolic KEGG GSEA confirmed OXPHOS as the most strongly upregulated pathway during ART initiation, with additional increases in TCA cycle, one-carbon metabolism, and fatty-acid metabolic pathways (Figure 3F-G). Finally, GSEA of DEGs comparing late ART (W23) with pre-infection (W0) revealed a broad downregulation of activation and metabolic pathways, accompanied by persistent disruption of cholesterol homeostasis (Figure. 3H). Metabolic KEGG GSEA further indicated sustained suppression of major energy pathways and a relative reliance on one-carbon metabolism and FA during late ART (Figure S9B).

Taken together, the metabolomics and RNA-seq data support our prior findings. To more clearly define the dominant metabolic alterations at peak infection and during ART, we integrated these datasets to construct a context-specific GEM of CD4⁺ T cells from SIV-infected macaques. Flux balance analysis was then applied to estimate the activity of individual metabolic reactions at each time point. This analysis revealed pronounced remodeling of lipid metabolism across all post-infection time points, along with an increase in positive flux through carbohydrate metabolic reactions following ART initiation (Figure 4A). Although lipid-associated reactions represented a greater proportion of total metabolic activity at all post-infection time points relative to pre-infection (Figure 4B), the fraction of anabolic lipid reactions (lipid synthesis) with positive flux decreased sharply—from nearly 50% pre-infection to ∼5% at W2 pi (Fisher’s exact test, adj *p*= 0.01; Figure 4C). These results provide an independent, quantitative validation of the fatty-acid synthesis shutdown identified by MIST during peak SIV replication. The only additional flux change that reached statistical significance in comparisons between pre- and post-infection time points was an increase in active transport reactions during late ART relative to pre-infection (BH-FDR adj *p*=0.03; Figure 4C).

**Fig. 4.**
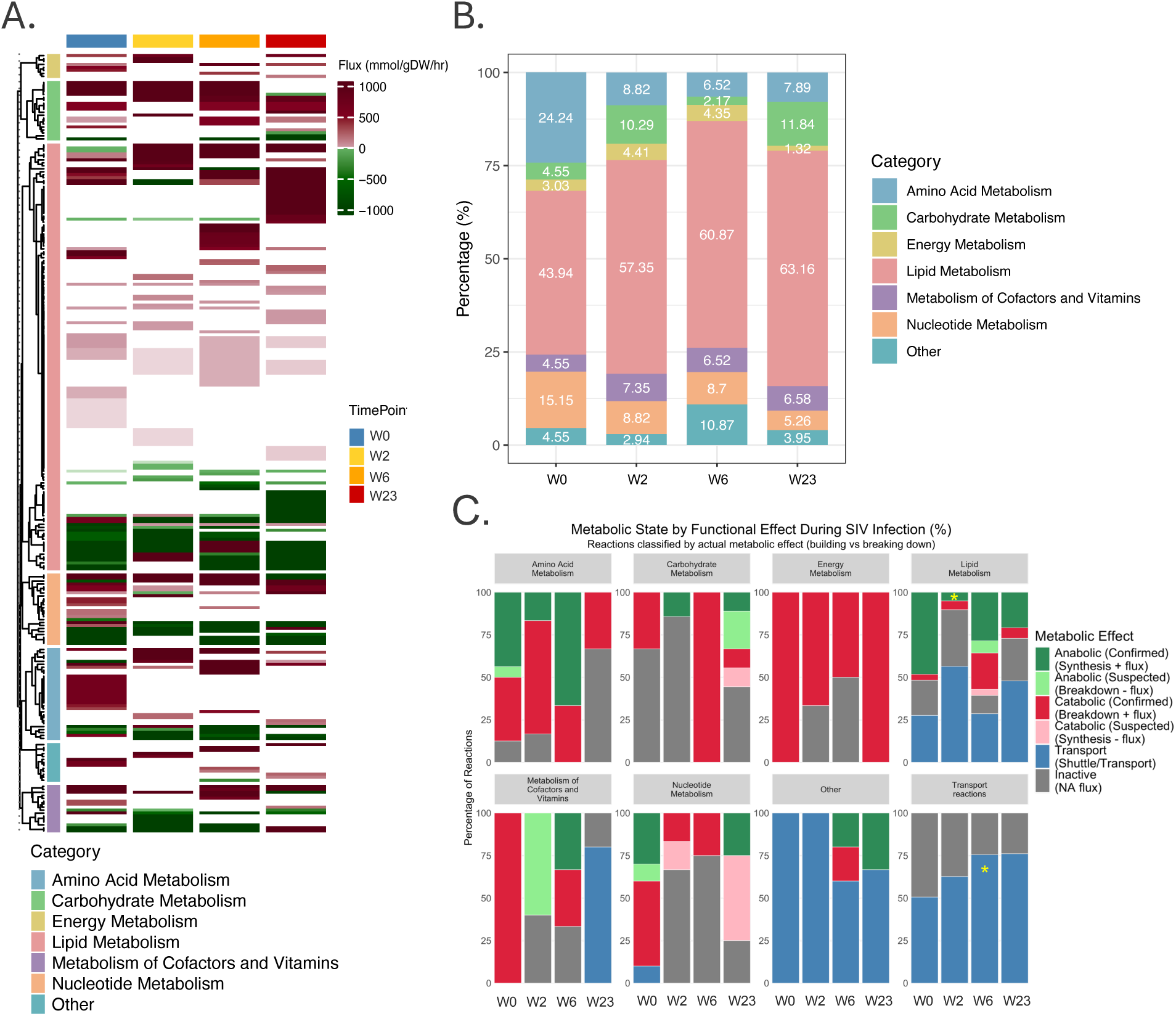
Genome Scale Metabolic Model (GEM) of CD4 T cells Confirms Shut Down of Lipid Anabolic Pathways at Peak Viral Replication. A-B) Contextualization of GEM specific to CD4⁺ T cells was performed using RNA-seq data from SIV-infected macaques across W0 (pre-infection), W2 (peak infection), W6 (week 3 post-ART), and W23. A) Heatmap of predicted flux values of all unique metabolic reactions is shown. The reactions are also annotated according to their metabolic super categories B) The stacked bar plots display the percentage of uniquely active metabolic reactions within major metabolic super-categories that were specific to each time point. C) Functional classification of GEM-predicted reaction fluxes was constructed after classification according to metabolic effect based on: (i) subsystem and reaction names, (ii) flux direction and (iii) *transport* or *shuttle* keywords. Frequencies of reaction types within each metabolic pathway were compared across time points using Fisher’s Exact test (* BH-FDR Adj p ≤0.05).

Given the profound impact of SIV infection on lipid metabolism in CD4⁺ T cells and the disruption of FA synthesis at peak infection identified above, we next asked whether these cellular metabolic changes were reflected in the circulating lipid profile. Plasma lipidomics revealed a progressive decline across all major lipid classes during acute SIV infection, including glycerolipids, glycerophospholipids, and sphingolipids (Figure 5A–B). Principal component analysis (PCA) showed that these lipid alterations accounted for most of the variance across infection and treatment stages, with PC1 and PC2 explaining 43.71% and 19.33% of the total variance, respectively (Figure 5C).

**Fig. 5.**
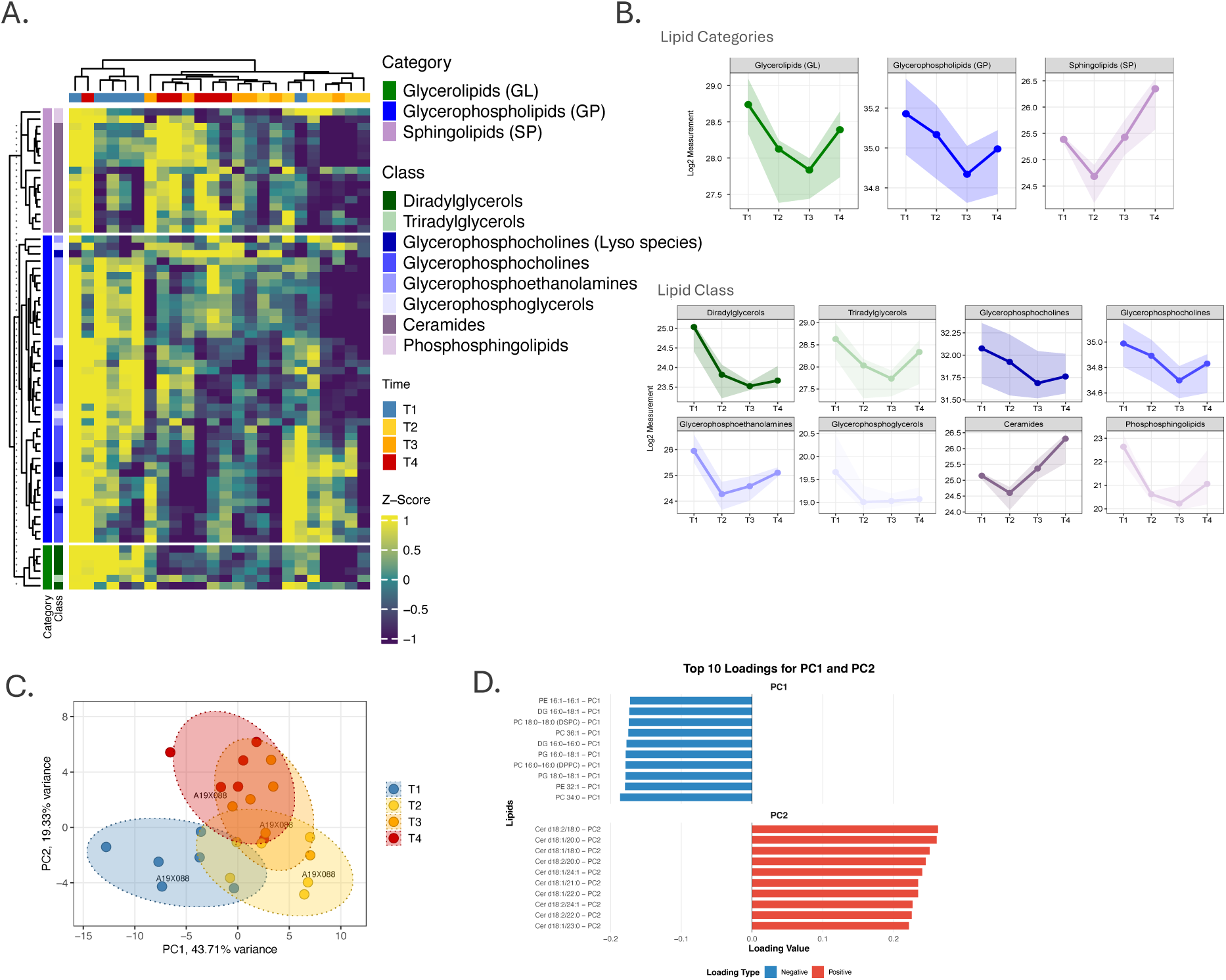
Plasma lipidomics confirms shut down of lipid synthesis at peak viral replication. A–B) Untargeted plasma lipidomics profiling was performed across T1 (pre-infection), T2 (peak infection), T3 (early ART), and T4 (late ART). A) Heatmap showing Z-scored abundances of major lipid categories and classes, including glycerolipids, glycerophospholipids, and sphingolipids. B) Line plots summarizing (median with lower (Q1) and upper quartiles (Q3) as shaded area) temporal changes in total lipid categories (top row) and individual lipid classes (bottom row) across time points. C) Principal component analysis (PCA) of plasma lipidomes across time points. Each point represents an individual sample, colored by time point, with 95% confidence ellipses shown. PC1 and PC2 together explain 63.0% of total variance. D) Loading plots identifying the top positive and negative contributors to PC1 and PC2.

PC1 primarily separated pre-infection and late ART time points (W0 and W23) from acute infection and early ART (W2 and W6), with the strongest loadings all driven by reductions in phospholipids and other structural lipid components, including phosphatidylcholines (PC), phosphatidylethanolamines (PE), phosphatidylglycerols (PG), and diacylglycerols (DG) (Figure 5D). These membrane lipids were markedly depleted during acute infection and remained suppressed through early ART, indicating substantial disruption of membrane biosynthesis. This pattern is consistent with our earlier MIST and metabolic modeling analyses and supports the conclusion that acute SIV infection induces a broad suppression of lipid synthesis, particularly affecting structural components of cellular membranes—an effect that is likely not restricted to CD4⁺ T cells. Notably, lipid classes displayed distinct recovery trajectories. Diacylglycerols, triacylglycerols, and glycerophosphoethanolamines exhibited incomplete recovery even during prolonged ART (Figure 5B). In contrast, sphingolipids—especially ceramides—declined sharply at peak viral replication (W2) but increased during ART, ultimately exceeding pre-infection levels by late ART (W23) (Figure 5A–B). Elevated ceramide species were the primary contributors to PC2, which best distinguished pre-infection and late ART time points from acute infection and early treatment stages (Figure 5D). This ART-associated increase in ceramides may have important implications for metabolic dysregulation during long-term therapy.

Together, these data demonstrate that acute SIV infection profoundly disrupts lipid homeostasis through a near-complete shutdown of FA synthesis, resulting in marked depletion of membrane phospholipids at peak viral replication. Although ART partially restores FA synthesis, the plasma lipid profile during late ART does not fully revert to the pre-infection state, with persistent deficits in specific lipid classes and compensatory elevations in others, most notably ceramides.

### OXPHOS levels during untreated infection associate with early ART reservoir size

To assess whether the metabolic changes observed in CD4⁺ T cells influenced viral replication or reservoir formation, we examined correlations between MIST-derived metabolic variables and cell-associated viral DNA (CA-vDNA) at defined time points. Reasoning that cellular metabolism during untreated infection may influence reservoir establishment, we correlated MIST variables measured at W2 with CA-vDNA levels during early ART. ATP5A levels (OXPHOS) in total CD4^+^ T cells at W2 positively correlated with CA-vDNA at W6, three weeks after ART initiation; however, this association did not remain significant after correction for multiple comparisons (p≤0.05; BH-FDR adj p>0.05; Figure 6A). Because CD4⁺ T cell subsets differ in susceptibility to infection and metabolic state, we performed a focused, hypothesis-driven subset-stratified analysis of ATP5A, testing whether W2 ATP5A levels in one or more memory CD4⁺ T cell subsets correlated more strongly with W6 CA-vDNA than ATP5A measured in bulk CD4⁺ T cells. Indeed, we found that ATP5A levels in EM cells strongly and significantly correlated with CA-vDNA at W6 post-infection, whereas correlations in other subsets were weaker or absent (Figure 6B). These data suggest that the bulk OXPHOS–reservoir association is concentrated in the EM compartment and support a model in which *ex*-OXPHOS-high EM cells during untreated infection may be preferentially represented in the reservoir after ART initiation.

**Fig. 6.**
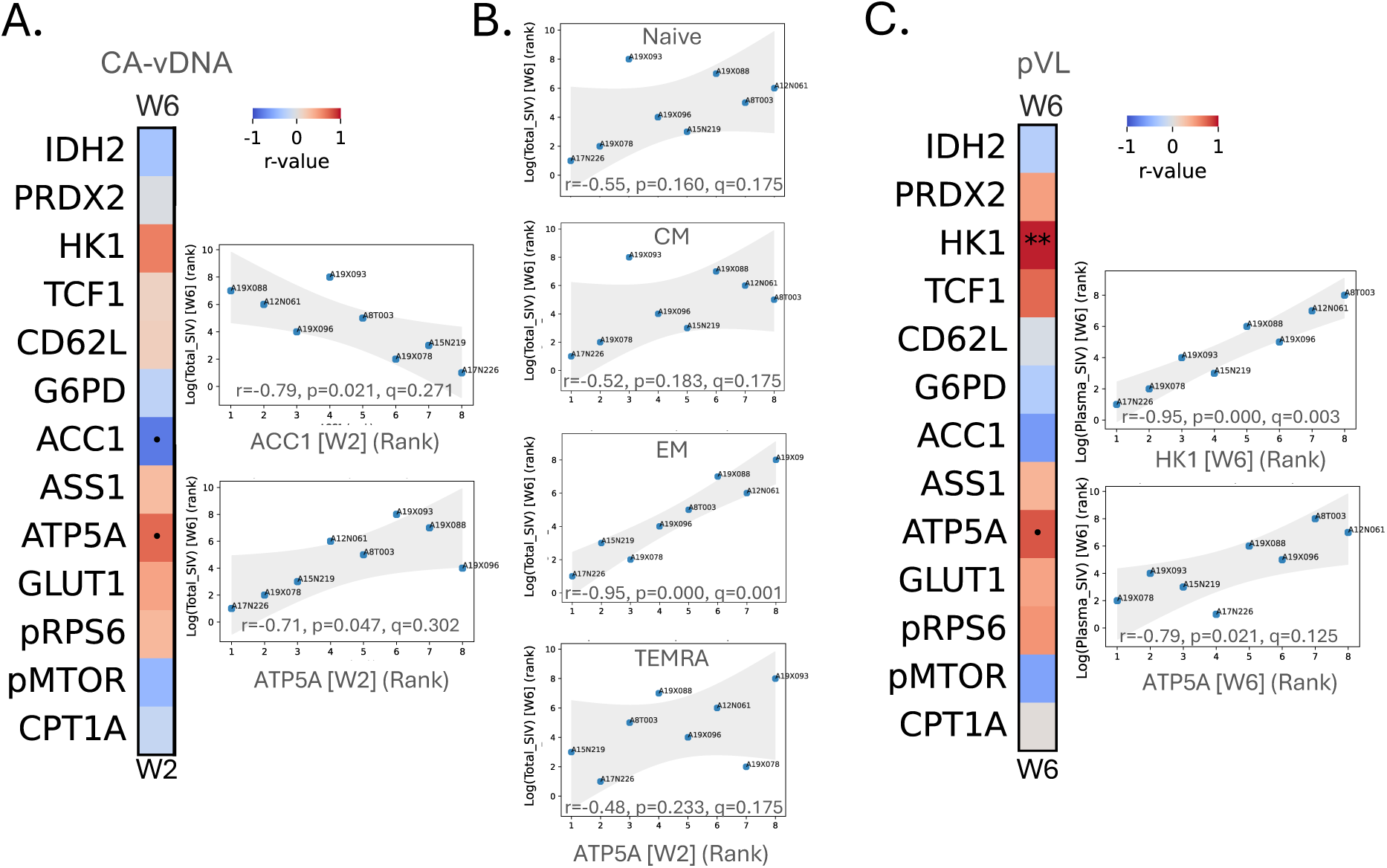
Virological variables correlations with MIST variables. A-B) MIST variables (MFI) at week 2 pi (peak) in bulk CD4^+^ T cells (A) and in CD4^+^ T cell subsets (B) were correlated with cell associated (CA) total vDNA at week 6pi C) MIST variables at W6 post infection were correlated with residual pVL at W6 post infection. A and C) Heatmap color is proportional to Spearman correlation coefficient. (*q≤0.05, **q≤0.01). Shown in dot plots are correlations with *p≤0.05 but q>0.05, where a single point did not drive the correlation (removal of any of the 8 samples did not impact the correlation).

In parallel, HK1 levels directly correlated with residual plasma viral load at W6, while ATP5A showed a positive trend (Figure. 6C; p≤0.05; BH-FDR adj p>0.05), suggesting that OXPHOS activity during ART initiation may also influence early reservoir transcriptional activity, whereas glycolytic flux appears to be a dominant driver of residual viral replication during early ART.

In contrast, ACC1 levels at peak infection tended to inversely correlate with both CA-vDNA and residual viral replication at ART initiation (Figure 6A–B). Combined with the marked reduction in ACC1 observed at W2, this pattern suggested that ACC1 downregulation was unlikely to be a direct consequence of viral replication within infected cells, but rather reflected antiviral responses in bystander CD4⁺ T cells.

### Type I IFN Responses Drive FA Synthesis Shut Down, Limiting Viral Replication

To directly test whether ACC1 downregulation reflected an antiviral bystander response rather than a cell-intrinsic consequence of productive infection, we performed in vitro HIV-1 infection experiments using MIST analysis. Infected CD4⁺ T cells (p24⁺ by intracellular staining) exhibited increased ACC1 expression compared to uninfected (p24⁻) cells within the same culture (Figure 7A; experimental conditions and gating strategy shown in Figure S10). These analyses, performed using an updated MIST2 panel (Table S4), further revealed downregulation of arginine and glutamine metabolism (ASS1 and GLS1) and OXPHOS in infected cells relative to uninfected cells within the infected culture. Importantly, when uninfected bystander CD4⁺ T cells (p24⁻) from infected cultures were compared to CD4⁺ T cells cultured in parallel under identical conditions but never exposed to HIV-1, ACC1 levels were significantly reduced in the bystander cells (Figure 7B). These results confirm that ACC1 downregulation is induced by the infectious environment rather than by productive viral replication.

**Fig. 7.**
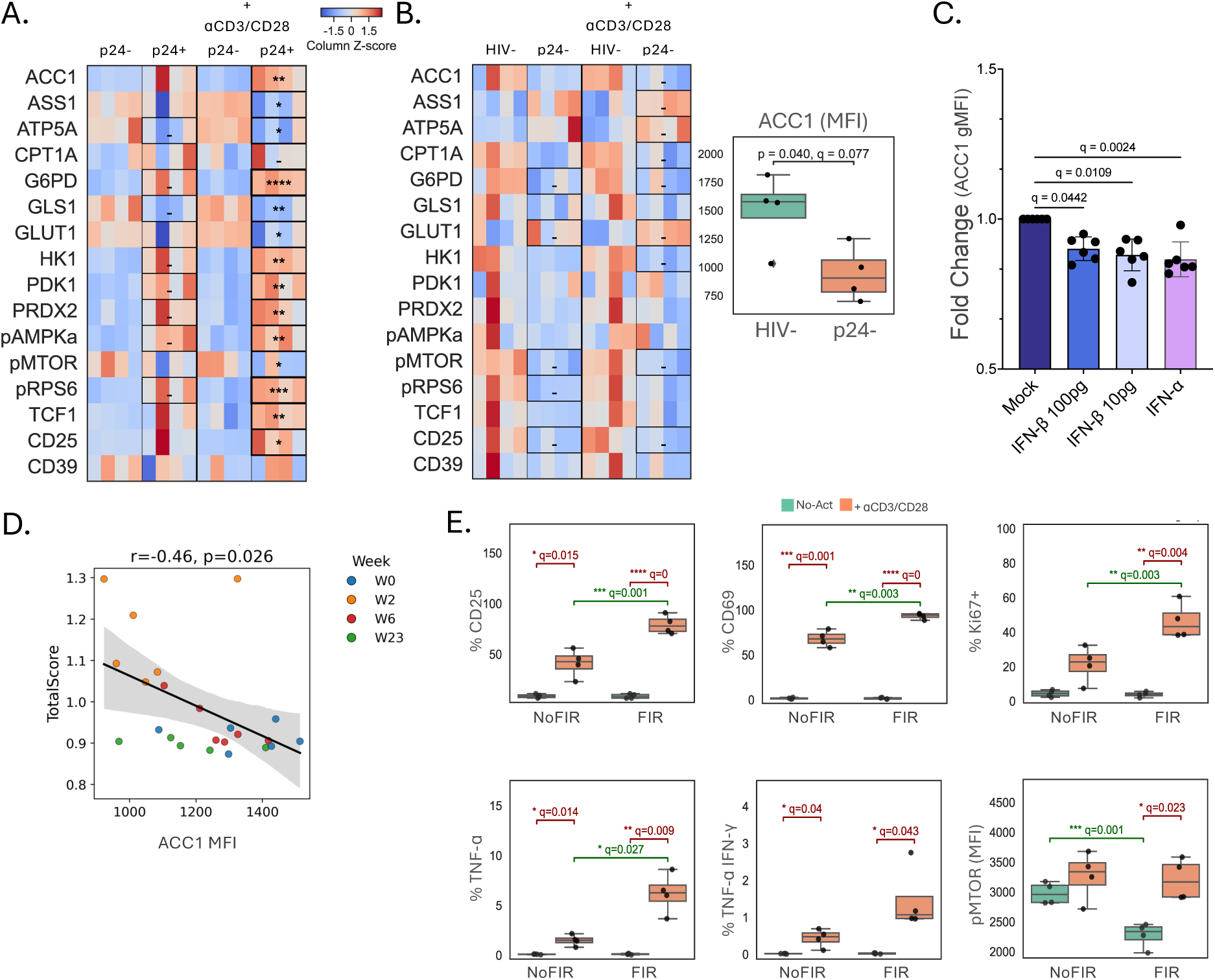
ACC1 Downregulation in bystander uninfected cells is driven by Type I IFN. A) Heatmap of MIST2 variables in p24^+^ CD4^+^ T cells compared to p24^-^ cells in the same culture in activated (anti-CD3/28) or non-activated cells. B) Heatmap of MIST2 variables in p24- cells in the HIV infected culture vs cultures never exposed to HIV (activated or non-activated cells). A-B) Paired two-sided t-tests with FDR-TSBH correction (-q≤0.10, *q≤0.05, **q≤0.01, ***q≤1e-3, ****q≤1e-4). Heatmaps display z-scored gMFI. Similar conclusions were obtained when the comparisons were done with non-parametric tests (see Supplementary Data File). C) Changes in ACC1 MFI levels after 24-48hrs of IFN-I exposure are shown (Friedman test with FDR-BH correction q values are shown). D) IFN transcriptional signature inversely correlated with ACC1 levels in SIV-infected macaque CD4^+^ T cells. Spearman rank correlation shown. E) Blocking ACC1 with Firsocostat (Fir) increases response to CD3/28 activation. Rhesus PBMC were activated anti-CD3/CD28 or left in culture in parallel without activation. The increase in frequency of CD25, CD69 and Ki67, cells producing TGF-α, TNF-α and IFN-γ and the levels of phospho-(activated)MTOR in activated (orange) compared non-activated (green) cells in presence and absence of Fir is shown compared with two-way repeated-measures ANOVA followed by paired t-test with FDR-TSBH correction (*q≤0.05, **q≤0.01, ***q≤1e-3, and ****q≤1e-4). Similar conclusions could be reached when the comparisons were done with non-parametric tests (see Supplementary Data File)

Previous studies suggested that type I interferon (IFN-I) signaling may suppress lipid biosynthesis in CD4⁺ T cells in a model of influenza virus infection^34^. Consistent with this, we found that treatment of human CD4⁺ T cells with IFN-I in vitro was sufficient to downregulate ACC1 expression (Figure 7C). Notably, higher IFN-I concentrations had diminishing effects on ACC1 suppression, suggesting a non-linear dose–response relationship. These findings indicate that the strong IFN-I responses observed at peak SIV infection (Figure 3) are likely key drivers of FA synthesis shutdown and lipid metabolic remodeling.

Supporting this conclusion, the IFN-I transcriptional signature, derived from RNA-seq module score analysis using canonical IFN-I–stimulated genes (ISGs; Table S5), inversely correlated with ACC1 protein levels measured by MIST across infection stages (Figure 7D). Interestingly, consistent with the in vitro dose–response data, the magnitude of ACC1 downregulation inversely, rather than directly, correlated with IFN-I signature strength at peak infection (Figure 7D).

ACC1 has been shown to regulate CD4⁺ T cell fate decisions, including the balance between regulatory T cells and Th17 cells^19^. Genetic deletion or pharmacological inhibition of ACC1 attenuates Th17-driven pathology^19^ and promotes Treg differentiation^35^, while reduced ACC1 expression has also been linked to enhanced memory potential in CD4⁺ T cells^18, 20^. To investigate the functional consequences of ACC1 downregulation during acute viral replication, we examined the effects of pharmacological ACC1 inhibition on T cell activation and HIV replication in vitro. Inhibition of ACC1 with Firsocostat (Fir, GS-0976) significantly enhanced CD4⁺ T cell responses to anti-CD3/CD28 stimulation (Figure 7E). Treated cells exhibited increased upregulation of activation markers CD25 and CD69, a higher frequency of Ki67⁺ proliferating cells, and increased production of TNF-α (Figure 7E). IFN-γ production also increased, although this did not reach statistical significance (Figure S11A). Notably, ACC1 inhibition reduced baseline phosphorylation of mTOR and RPS6 in resting cells; however, upon TCR stimulation, p-mTOR levels increased to those observed in untreated cells (Figure 7E and Figure S11A). Similar effects were observed in human PBMCs (Figure S11B). Together, these data indicate that direct suppression of FA synthesis enhances TCR responsiveness, consistent with a heightened antiviral effector state.

Finally, we evaluated the direct impact of ACC1 inhibition on HIV-1 replication. Consistent with previous reports showing that the ACC1 inhibitor soraphen A suppresses HIV-1 replication in vitro^36^, inhibition of ACC1 with Firsocostat similarly reduced HIV-1 replication in primary CD4⁺ T cells (Figure S12). Collectively, these findings demonstrate that at peak SIV replication, IFN-I responses drive a profound reprogramming of lipid metabolism characterized by shutdown of de novo FA synthesis. Independent of IFN-I, FA shut-down enhances T cell effector responsiveness while simultaneously limiting viral replication, indicating that suppression of FA synthesis represents both an antiviral host defense mechanism and a direct constraint on HIV replication.

## Discussion

Acute HIV/SIV infection and the early period surrounding ART initiation are critical phases that shape viral replication, immune responses, and the establishment of the long-lived viral reservoir^37, 38^. These early events are also known to influence virologic control following analytical treatment interruption (ATI), and thus the likelihood of achieving durable remission or cure^39, 40^. Because cellular metabolism regulates susceptibility to infection, viral transcription, and T cell antiviral responses^5, 41^, defining how CD4⁺ T cell metabolism is remodeled during these phases is essential for understanding mechanisms of viral persistence. Here, by integrating MIST profiling, transcriptomics, lipidomics, genome-scale metabolic modeling, and functional assays, we identify IFN-driven shutdown of de novo FA synthesis as a novel antiviral effector mechanism during acute HIV/SIV infection, a finding with direct functional consequences for T cell responses and viral replication. We further characterize a second, distinct metabolic phase during ART, defined by broad metabolic suppression with selective preservation of OXPHOS, and identify a correlative link between early OXPHOS levels and reservoir establishment.

The most prominent metabolic change during acute infection was a strong inhibition of FA synthesis, converging across all platforms: ACC1 protein levels by MIST, reduced FA-dependent respiration by Seahorse, membrane phospholipid depletion by lipidomics, and near-complete suppression of anabolic lipid reactions in the GEM. Prior studies based primarily on HIV-infected cell lines reported that HIV replication increases FA synthesis and upregulates FASN to support virion assembly, and that inhibition of FASN limits viral production^16^. Our findings do not contradict this; instead, they suggest that those earlier observations may apply specifically to the relatively rare, infected cells undergoing active viral replication. In contrast, the majority of CD4⁺ T cells in vivo during acute infection show a coordinated suppression of FA synthesis that is particularly pronounced in memory subsets.

We found that FA-synthesis shutdown during acute infection was closely linked to IFN-I responses. Strong ISG signatures were present at peak infection, IFN-I treatment of human and rhesus CD4⁺ T cells reduced ACC1 expression in vitro, and ACC1 levels inversely correlated with the IFN-I transcriptional score across infection stages. Importantly, pharmacologic ACC1 inhibition with Firsocostat significantly enhanced CD4⁺ T cell responses to TCR stimulation and directly suppressed HIV replication in vitro, even in the absence of IFN-I signaling. This indicates that FA-synthesis shutdown represents an antiviral mechanism that operates independently of, and downstream from, IFN-I. These observations add to reports in other viral infections of IFN-I-driven suppression of lipogenic enzymes, including FASN, as part of an antiviral program^17, 18^ and directly challenge the prevailing view that ACC1 suppression and FA-synthesis inhibition primarily serves to promote memory T cell formation^20^ or a regulatory T cell phenotype^19^. Here, in the context of acute viral infection, ACC1 inhibition enhances effector T cell responses and limits viral replication, defining a novel antiviral function for lipid synthesis suppression.

A small paradoxical finding was that animals with the highest ISG expression tended to have less ACC1 downregulation. Although the underlying reason is unclear, this was supported by in vitro data showing smaller effect with increasing levels of IFN-I and may reflect differences in timing of ACC1 turnover, saturation of upstream signaling intermediates, or compensatory pathways. Overall, our data warrant further exploration of the impact of IFN-I on lipid metabolism.

In addition to FA synthesis, acute infection affected other metabolic pathways. Both metabolomics and the GEM model indicated disruption of the TCA cycle, altered glutamate–αKG balance, and changes in multiple transport reactions. These broader metabolic alterations are consistent with high inflammatory signaling, cellular stress, and potentially increased apoptosis and cellular turnover during acute infection. Taken together, the data show that acute infection imposes a coordinated metabolic state characterized by suppression of anabolic lipid metabolism and mitochondrial remodeling.

The transition into ART initiation marked a second, distinct metabolic phase. Although viral replication was rapidly suppressed, CD4⁺ T cell metabolism did not revert to the pre-infection state. Instead, we observed a broad suppression of major metabolic pathways, including glycolysis, FAO, redox balance, one-carbon metabolism, and other anabolic processes.

This global contraction most likely reflects reduced antigenic and inflammatory stimulation following viral suppression, rather than direct pharmacological effects of antiretroviral drugs. Supporting this interpretation, previous data^42^ and our own work (Figure S13) show that the antiretroviral classes used in our regimen (NRTIs, INSTIs) reduce rather than preserve mitochondrial function in primary T cells. Our data suggest that these alterations may emerge early during ART initiation as a consequence of early metabolic reprogramming, and may have important functional consequences for T cell immunity^43, 44^. Consistent with this possibility, individuals on long-term ART display persistent metabolic defects, including reduced metabolic flexibility and impaired T cell activation^45, 46^.

Within this overall metabolic shutdown, OXPHOS emerged as a notable exception. Subset-stratified analysis revealed that the early-ART OXPHOS increase was most pronounced in memory CD4⁺ T cell subsets, whereas by W15 the elevation extended to all subsets including naïve cells. Elevated OXPHOS has previously been associated with adverse clinical outcomes, including higher viral set points and impaired CD4⁺ T cell recovery^21, 47^. Our data indicate that this sustained reliance on mitochondrial respiration begins early during ART, coinciding with the period when long-lived infected cells are thought to transition into the latent reservoir. Importantly, this metabolic pattern observed in the SIV macaque model was recapitulated in PWH, supporting the translational relevance of our findings.

Given the central role of mitochondrial respiration in supporting the effector-to-memory transition and the establishment of quiescent CD4⁺ T cell states^48^, elevated OXPHOS in effector CD4 T cells during untreated infection may represent a metabolic driver of latency establishment and reservoir persistence in infected cells transitioning to a memory, quiescent state. This interpretation remains correlative and will require direct lineage or fate-mapping approaches to establish causality.

Several limitations of this work should be considered. This study was performed in a relatively small macaque cohort, with most analyses derived from 6–8 SIV-infected female macaques and relied on a single infection model (SIVmac239) with early ART initiation. Hence, it may not fully capture the metabolic trajectories associated with mucosal HIV acquisition or the more prolonged untreated viremia in PWH. Given documented sex-based differences in immune responses and metabolism^49, 50^, generalizability of these findings to male subjects requires confirmation in future studies. In addition, our analyses focused on circulating CD4⁺ T cells, whereas metabolic responses to infection and viral suppression may differ across tissues and immune cell types, particularly within lymphoid and mucosal sites that are central to reservoir establishment. Replication of these findings in independent male macaque cohorts, extension to tissue-resident cells, and in vivo studies with additional viruses will be important to determine the extent to which IFN-I–driven FA synthesis shutdown represents a conserved antiviral mechanism.

Finally, because of the extremely low frequency of productively infected cells (<0.1% of CD4⁺ T cells), we could not analyze metabolic changes specifically within infected CD4⁺ T cells in vivo. Therefore, our findings primarily reflect metabolic changes within the overall circulating CD4⁺ T cell population, which is overwhelmingly composed of uninfected or latently infected cells. Future studies with substantially larger numbers of infected-cell events will be required to robustly define metabolic reprogramming within productively infected CD4⁺ T cells.

Taken together, our data define a temporal sequence of metabolic states that shape early SIV infection. Acute infection is characterized by a strong IFN-linked suppression of FA synthesis that enhances antiviral T cell responses and directly limits viral replication — a novel effector mechanism that reframes the functional significance of lipid metabolic reprogramming during acute viral infection. ART induces a second, distinct phase of broad metabolic contraction with selective preservation of mitochondrial OXPHOS, the levels of which correlate with early reservoir formation. These distinct metabolic phases highlight potential windows for intervention: transient reinforcement of FA synthesis inhibition during acute infection could strengthen antiviral defenses and limit early reservoir seeding, while targeting OXPHOS at ART initiation could potentially interfere with reservoir establishment.

In summary, our findings demonstrate that host metabolic programs actively shape both untreated and treated HIV/SIV infection. By defining stage-specific metabolic adaptations and identifying IFN-driven FA synthesis suppression as a bona fide, novel antiviral effector mechanism, this work opens new avenues for metabolic strategies aimed at limiting viral persistence and supporting durable immune control.

## Methods

### Study design and Ethics Statement

A total of 8 adult female Indian origin *Rhesus* macaques (*Macaca mulatta*; Mamu A*01, B*08 and B*17 negative) were used for the study described in this manuscript (Fig. 1A). All macaques were selected form the colonies bred and raised at the New Iberia Research Center (NIRC), University of Louisiana at Lafayette. All animal experiments were conducted following guidelines established by the Animal Welfare Act and the NIH for housing and care of laboratory animals and performed in accordance with institutional regulations after review and approval by the Institutional Animal Care and Usage Committees (IACUC) of the University of Louisiana at Lafayette (2023-003-8841; protocol 8841-02). Rhesus macaques (n=8 main study + 3 separate study) were infected with 300 TCID_50_ of the barcoded SIVmac239M2 stock intravenously and ART (Tenofovir [PMPA] at 20mg/ml, Emtricitabine [FTC] at 40mg/ml and Dolutegravir [DTG] at 2.5mg/ml) was initiated on week 3 pi. Blood viral load was monitored biweekly before and during ART and PBMC were collected at indicated time points (Figure 1A).

### Plasma and Tissue SIV Viral loads (VL)

Blood was collected in EDTA tubes and plasma was separated by centrifugation and used for the determination of plasma VL by SIVgag qRT-PCR at NIRC or at Leidos (Quantitative Molecular Diagnostics Core, AIDS and Cancer Virus Program Frederick National Laboratory). PBMCs were separated through Ficoll density gradients, washed counted and cryopreserved. Cell associated (CA)-vDNA levels were assessed in DNA extracted from snap frozen PBMC pellets by SIV IPDA^51^ with the following modifications: CCR5 was used as a host genomic control instead of RPP30, SIV pol and env were as published, the assay was performed on the Applied Biosystems™ QuantStudio™ Absolute Q™ Digital PCR System (Thermo Fisher Scientific) with amplification step set at 30 seconds. CCR5 primers and probes were as following: (exon 1, 5’-3’) Fw CCAGAAGAGCTGCGACATCC; Rv GTTAAGGCTTTTACTCATCTCAGAAGCTAAC; Probe: TTCCCCTACAAGAAACTCTCCCCGGTAAGTA; (Exon 2, ∼2,000bp after Exon 1; 5’-3’) Fw TCACAGGGTGGAACAAGATG; Rv TTATACATCGGAACCCTGCC and Probe: CAAGTGTCAAGTCCAACCTATGACATCGA. We used a multiplex set up with the following fluorescence channels: FAM for SIV pol, VIC for SIV env, and ABY and JUN for the two CCR5-based host targets. Primer and probe concentrations were optimized and used at final concentrations of 600 nM and 200 nM, respectively for all sets. For each reaction, 300 ng of genomic DNA was analyzed using the Absolute Q DNA Digital PCR Master Mix in a final reaction volume of 10 µL. Data acquisition and absolute quantification were performed using the Absolute Q analysis software. Absolute target concentrations were derived from Poisson estimates based on the fraction of positive partitions and adjusted for sample dilution. Cell equivalents were estimated from CCR5 copy numbers assuming two alleles per diploid genome using the higher of the two CCR5-derived values. Intact viral copies were estimated by normalizing the number of double positive pol and env copies by the shearing index calculated from the duplex CCR5 control. Final CA-vDNA values were expressed as copies per 10⁶ cell equivalents.

### Metabolic Immune Signature Technology (MIST) Assay

PBMC from macaques at indicated time points were thawed in AIM V media (Gibco) with 50U/ml of Benzonase. Cells were washed and stained with the fixable viability dye FVS440UV. Following a wash and Fc receptor blocking, surface staining was performed in FACS buffer 30 minutes at 4 °C. Antibodies for MIST1 are listed in Table S1 and for MIST2 in Table S4. Cells were then washed twice, fixed and permeabilized using the Invitrogen eBioscience Fix/Perm Buffer for 30 minutes at 4 °C. Intracellular staining was performed in Perm buffer supplemented with CellBlox Plus Blocking Buffer and mouse IgGs for 45 minutes at room temperature (RT). Stained cells were acquired immediately on a Cytek Aurora 4-laser spectral flow cytometer. Spectral unmixing and spillover correction were performed using SpectroFlo software. Downstream gating and analysis of total CD4⁺ T cells, CD4⁺ T cell subsets, and CD4⁻ T cells were conducted on fully compensated and quality-controlled data, with metabolic marker levels quantified as geometric mean fluorescence intensity (gMFI). The MIST assay was validated using isolated CD4⁺ T cells and PBMC from naïve rhesus macaques cultured in RPMI 10% FBS with Glutamax at 1.5M cells /mL either unstimulated (with 20U/ml of IL-2) or stimulated for 3 days on anti-CD3 (FN18 Nonhuman Primate Reagent Resource, NHPRR) and anti-CD28 (28.8, NHPRR) coated plates with 50U/mL of IL2. Following stimulation, cells were harvested and processed as above.

### RNA-seq of sorted CD4^+^ T cells and RT- dPCR for ACC1

Cryopreserved PBMCs from weeks -1, 2, 6, and 23 were stained with CD3 (BD; #566955; SP34-2) and CD8 (BioLegend; #344706; SK1) surface antibodies. CD8 negative T cells were sorted on a BD FACSAria-5 laser cell sorter (Flow Cytometry Core, Robert H. Lurie Comprehensive Cancer Center) at a flow rate of 1. Sorted cells were allocated for Agilent Seahorse Assay and remaining cells were pelleted and stored at -80C until processing. Total RNA was extracted using the QIAGEN RNeasy Mirco Kit with on-column DNase treatment. RNA concentration and quality were assessed by high-sensitivity RNA electrophoresis at the Northwestern University Metabolomics Core Genomics Branch. 10ng of total RNA per sample was submitted to the core for metabolic mRNA library preparation, followed by sequencing using NextSeq 2000 P2 flow cell pair-end reads resulting in >20M reads per sample. Samples from 3 macaques (A15N219, A19X078, A19X093) with enough left-over RNA were assayed also by digital PCR. RNA was converted to cDNA using the SuperScript Vilo Kit (Thermo Fisher) and cDNA (5ng equivalent of RNA) was run on the Absolute Q dPCR platform using: Forward: GAAGAGGCACCAGCTACTATTG Reverse: CAGTCCCAGCACTCACATAAC; Probe: /56-FAM/CAGTGTGCG/ZEN/GTGAAACTTGCCAAA/3IABkFQ/

### RNA-seq Analysis

RNA sequencing data were processed using the nf-core/rnaseq workflow (v3.19.0; doi: 10.5281/zenodo.1400710), part of the nf-core community-curated pipeline collection^52^. The workflow was executed with Nextflow v25.04.4^53^ and leveraged reproducible software environments provided by Bioconda^54^ and BioContainers^55^. The Macaca mulatta reference genome (Mmul 10) and corresponding gene annotation files were obtained from Ensembl and used as the reference for all RNA-seq analyses. Read alignment and transcript quantification were performed within the nf-core/rnaseq pipeline using the STAR–Salmon mode. In this configuration, STAR^56^ was employed as the primary splice-aware aligner to map sequencing reads to the reference genome. Following alignment, transcript abundance estimation and gene-level quantification were carried out using Salmon in alignment-based mode^57^ leveraging STAR-generated alignments. This approach combines the high alignment accuracy of STAR with the robust transcript-level quantification capabilities of Salmon, including correction for sequence-specific and GC-content biases. Gene-level expression estimates were derived by aggregating transcript-level abundances according to the provided Ensembl gene annotation. Gene-level count matrices generated by the pipeline were used for all downstream analyses.

Differential gene expression analysis was conducted using DESeq2 v1.44.0. A likelihood ratio test (LRT) framework was applied to identify genes exhibiting significant temporal expression changes across time points. To account for unwanted technical variation, factors of unwanted variation were estimated using the RUVSeq package v1.38.0. Residual-based estimation was employed, and the inferred unwanted variation factor was incorporated into the DESeq2 design matrix to mitigate potential bias arising from unmodeled technical effects. The final statistical model included subject specific effects, the estimated unwanted variation factor, and time as explanatory variables. The reduced model for the LRT excluded the time variable, enabling identification of genes with significant time-dependent expression changes.

Time-series clustering of gene expression profiles was performed using TMixClust v1.26.0 (DOI: 10.18129/B9.bioc.TMixClust). Only genes identified as significantly time-dependent in the LRT analysis (adj p≤0.05) were included. Transcript abundance values (TPM) were z-score normalized, and average z-scores for each time point were used as input for clustering. The optimal number of clusters was determined based on silhouette width. Gene set enrichment analysis (GSEA) was carried out using the enrichr module implemented in gseapy v1.1.5^58^ ^59^ to identify biological pathways associated with each temporal expression cluster.

GSEA was performed on all protein-coding genes ranked by log2 fold change using the fgsea (v1.32.2). A preranked enrichment approach was applied to evaluate pathway-level transcriptional changes of each time point against the other. Hallmark gene sets were obtained from the MSigDB. KEGG pathway gene sets were retrieved from the C2 curated gene set collection (CP: KEGG subcategory) using the msigdbr (v7.5.1). In addition to global pathway analysis, a curated subset of 66 metabolism-associated KEGG pathways was manually selected for focused enrichment analysis (Tables S2 and S3). Pathway enrichment was evaluated using normalized enrichment scores (NES) and multiple-testing–adjusted *P* values, with positive NES values indicating enrichment among upregulated genes and negative NES values indicating enrichment among downregulated genes.

To correlate IFN gene signature with the level of ACC1 expression, Type I IFN transcriptional activity scores were calculated using a rank-based single-sample gene set scoring approach. equivalent to the R package singscore, implemented in Python^60^. Scores were derived from median-of-ratios–normalized RNA-seq data of IFN-I gene set (Table S5) and scaled to theoretical rank bounds, yielding values between 0 and 1.

Bulk RNA-seq data were normalized using the median-of-ratios method, and the resulting normalized expression matrix was used as input for scoring. Gene-wise ranks were computed independently for each sample using the rankGenes() function, converting absolute expression values into relative within-sample ranks. An interferon activity score was then calculated for each sample using uncentered scoring to preserve raw rank-based enrichment values. The resulting score represents a continuous measure of coordinated interferon pathway activation, with higher scores indicating stronger induction of the interferon response program.

### Plasma Metabolomics and Lipidomics

Plasma samples were processed for hydrophilic metabolomics and lipidomics using methanol-and a methyl tert-butyl ether (MTBE-) based extraction methods, respectively. For metabolite extraction, clarified plasma was mixed with ice-cold methanol at 4:1 (vol/vol) ratio, vortexed intermittently, and incubated at -20°C to precipitate proteins. Samples were centrifuged at high speed, and the metabolite-containing supernatant was collected. For lipidomics, plasma was combined with a methanol:MTBE (3:10, vol/vol) solution at a 5:1 ratio, vortexed, incubated at - 20°C, and centrifuged to induce phase separation. The organic, lipid-containing phase was collected and stored at –80°C until transfer to the Northwestern Metabolomics Core Facility for Comprehensive Hydrophilic Metabolites Panel (CHMP) using Liquid Chromatography Mass Spectrometry (LC-MS).

For metabolomics and lipidomic analysis, total ion content (TIC) normalized metabolites intensities were first log2-transformed after adding a pseudocount of 1 and used for all downstream analyses. Temporal differences in metabolite abundance were assessed using the Friedman test implemented in R. Time-series clustering of metabolite profiles was performed using TMixClust v1.26.0 (DOI: 10.18129/B9.bioc.TMixClust). Log2-transformed data were z-score normalized, and average z-scores across time points were used as clustering input. The optimal number of clusters was determined using silhouette width.

TIC normalized lipid intensities were first log2-transformed with the addition of a pseudocount of 1 prior to analysis. Principal component analysis (PCA) was performed using PCAtools v2.16.0 to explore global variation and temporal patterns in lipid abundance across samples.

### Agilent Seahorse Assay

Sorted CD4^+^ T cells (negative selection, CD8^-^ T cells) were resuspended in Agilent Seahorse XF RPMI media supplemented with 0.1-0.2mM glucose, 1mM pyruvate, and 2mM glutamine and seeded at 90,000 cells per well of an HS Mini SeaHorse plate in duplicate or triplicate (depending on sorted cell availability), and subjected to sequential injections of Etomoxir (4 mM), Oligomycin (1.5mM), BAM15 (2.5mM), and Rotenone/Antimycin A (0.5mM). The HS Mini plate and sensor cartridge were loaded into an Agilent HS Mini Analyzer and ran using the manufacturer’s T Cell Metabolic Profiling with Acute Injection protocol. The data were analyzed using the Agilent SeaHorse Analytics software. The etomoxir induced inhibition of OCR was calculated by comparing the mean of the three pre- and post-etomoxir OCR readings and expressed as a percentage of mean basal OCR.

### Genome Scale Metabolic Model (GEM)

GEM for Macaca mulatta was generated following the pipeline described by Wang, Hao et al^61^. The model was constructed using the Human1 GEM^62^ as a template, with orthologous gene mappings between human and Macaca mulatta obtained from Ensembl. The updateAnimalGEM function from the RAVEN2 package^63^ was employed to adapt the human model to Macaca mulatta, incorporating species-specific metabolic pathways and reactions.

To ensure metabolic completeness, gap filling was performed using the gapfill4essentialTasks function, enabling the reconstructed GEM to carry out essential metabolic functions and biomass production. The curated Macaca mulatta GEM was subsequently used to generate a context-specific model representing SIV-infected conditions. Context-specific model reconstruction was performed using the Fast Task-driven Integrative Network Inference for Tissues (ftINIT) algorithm^64^. TPM-normalized transcriptomics data served as input, with a minimum expression threshold of TPM ≥1 applied to define active genes. The resulting context-specific GEM was subjected to flux balance analysis (FBA), using ATP hydrolysis as the objective function. FBA was conducted using the solveLP function implemented in the RAVEN toolbox v2.10.1^63^. To summarize functional metabolic states from FBA and compare the proportion of anabolic, catabolic, transport, and inactive reactions across time points, reactions predicted by the context-specific GEM were classified using a rule-based algorithm implemented in R. Reactions with missing, non-numeric, or zero flux were classified as inactive. Transport reactions were identified based on the terms *“transport”* or *“shuttle”* in reaction names or subsystem annotations. Non-transport reactions were classified as anabolic or catabolic based on reaction names or subsystem annotations as anabolic (including biosynthesis, synthesis, elongation, desaturation, formation, transferase, methyltransferase, ligase, synthetase, *and* reductase) or catabolic (including oxidation, degradation, hydrolysis, catabolism, breakdown, hydrolase, peptidase, dehydrogenase, *and* oxidoreductase). Flux direction was used to confirm or invert functional classification, with positive flux indicating pathway engagement and negative flux interpreted as the opposite functional effect. When explicit annotations were absent, reactions were classified based on assigned super-category (e.g., amino acid or carbohydrate metabolism).

### Single Cell Human GEM

Published scRNA-seq datasets from NCBI BioProjects PRJNA662927 and PRJNA681021 were used to study the metabolic modulation in human CD4⁺ T cells from HIV-negative (n = 4), ART-experienced (n = 6), and treatment-naïve individuals (n=6)^32, 33^. Raw FASTQ files were processed using Cell Ranger^65^, and downstream analyses such as pre-processing, normalization, and batch correction were performed with the Seurat v5.0.0 R package^66^. Gene expression matrices corresponding to CD4⁺ naïve and CD4⁺ central memory T cells were extracted and aggregated within each cohort to generate pseudo-bulk profiles. The aggregated expression data were normalized using counts per million (CPM). These normalized matrices were used for context-specific genome-scale metabolic modeling and flux balance analysis as described above, and human generic GEM was used as the reference.

### In vitro HIV Infection and Parallel Uninfected Culture MIST analysis

PBMCs were cultured in RPMI supplemented with 10% FBS and GlutaMAX, with IL-2 (20 U/mL for non-activated or 50 U/mL for activated conditions), on anti-CD3/CD28–coated plates (activated) or mock-coated plates (non-activated) for 48 hours. Cells were then either harvested and stained with the MIST panel to profile metabolism in HIV-unexposed CD4⁺ T cells or infected with HIVBal. Infection was performed via spinoculation with 3 x 10^4^ TCID50 of HIV-Bal per 1M cells for 2hrs at 2000rpm, 37 °C. The cells cultured for an additional 7 days. Infected cultures were stained with the MIST panel together with intracellular HIV p24. Productively infected and uninfected CD4⁺ T cells were identified based on p24 expression (p24⁺ and p24⁻; see staining protocol in the next paragraph), and geometric mean fluorescence intensities (gMFI) of metabolic markers were quantified. Paired analyses were performed separately for activated and non-activated conditions to distinguish infection-specific metabolic reprogramming (p24⁺ vs p24⁻ cells within infected cultures) from bystander or infection-environment–driven effects (p24⁻ cells from infected cultures vs CD4⁺ T cells from uninfected cultures under the same conditions for the same number of days). HIV-Bal stock was prepared by infection of activated human PBMC using a previous stock and titrated on TZM-bl cells.

### Firsocostat (GS-0976) Impact on CD4^+^ T cell activation and HIV infection

To determine the impact of ACC1 inhibition on T cell activation, PBMCs from three naive rhesus macaques and four human donors were thawed and cultured in Human Plasma–Like Medium supplemented with dialyzed fetal bovine serum, GlutaMAX, and IL-2 at 1.5 × 10⁶ cells/ml with and without Firsocostat (500 nM) under non-activated or activated conditions, with activation achieved by culturing on anti-CD3/CD28 coated plates. Non-activated cultures were maintained in medium containing IL-2 at 20 U/ml, whereas activated cultures were cultured in the same medium supplemented with IL-2 at 50 U/ml. Firsocostat was maintained throughout the activation period, and cells were cultured for 2 days with Brefeldin A added 5 hours before harvest. Cells were stained for flow cytometric analysis using the FoxP3 Transcription Factor Staining Buffer Set (Invitrogen) together with an activation and metabolic panel including CD3, CD4, CD25, CD69, TNF-α, IFN-γ, Ki-67, phosphorylated mTOR (Ser2448), and phosphorylated RPS6 (Ser235/236). Frequencies of activation markers as well as median fluorescence intensities of phosphorylated metabolic markers were quantified on gated CD3⁺CD4⁺ T cells using a uniform gating strategy across all donors and conditions.

To determine the impact of ACC1 inhibition on in vitro HIV infection of human CD4^+^ T cells, isolated CD4^+^ T were thawed and activated for 72 hours. Activated cells were infected with Bal-HIV via spinoculation and, following infection, the cells were treated with Firsocostat (GS-0976) at 500 nM, 50 nM, or vehicle control in RPMI 10% FBS with 20U/ml of IL2. After 24 hours, the virus was washed from the suspension, and the cells were replated in fresh media with 20U/ml IL-2 and corresponding treatment conditions. Percentage of p24+ cells was acquired on days 4 and 7 post infection by intracellular staining following viability dye labeling, fixation and permeabilization with the BD Cytofix/Cytoperm Kit, and staining with anti-HIV-1 p24 antibody (KC57, NIH HIV Reagent Program, Division of AIDS, NIAID, NIH, ARP-13449, contributed by DAIDS/NIAID) and acquired via Cytek Aurora spectral flow cytometer. Cell pellets and supernatants were collected at defined timepoints and stored at -80°C for downstream RT-qPCR analysis using gag/CCR5 multiplex primers and probes as in ^65^ and a Thermo Fisher QS3 with default cycling conditions

### In vitro ART Impact on Rhesus PBMC

Uninfected Rhesus PBMC from 5 naïve macaques were thawed and cultured in R10 and 10U/ml of IL-2 for 7 days at 1.5 million cells/ml with or without Tenofovir (TDF, 1μM; FTC 5μM; DTG 5μM) added to the culture every other day. On day 8^th^, cells were collected and stained for MIST analysis.

### Statistics and Reproducibility

Statistical analyses were performed using R (version 4.3) and Python (version 3.9). All tests were two-sided. Multiple testing correction was applied as indicated for each analysis using the two-stage Benjamini–Hochberg false discovery rate method (TSBH-FDR). Adjusted *q* values are reported throughout. For MIST panel validation and in vitro activation experiments, paired comparisons between activated (anti-CD3/CD28) and non-activated conditions were performed within matched samples using Wilcoxon signed-rank tests, with TSBH-FDR correction applied across metabolic variables. These analyses were conducted using gMFI values within total CD4⁺ T cells as well as within defined CD4⁺ T cell subsets. For in vitro infection experiments with small sample sizes (n = 4), paired two-sided *t* tests were used to compare p24⁺ and p24⁻ CD4⁺ T cells within the same infected culture, as well as to compare p24⁻ CD4⁺ T cells from infected cultures with HIV⁻ CD4⁺ T cells from matched uninfected control cultures processed in parallel. Non-parametric test results (Wilcoxon signed-rank t tests results) are also reported in the Supplementary Data File 1. Longitudinal time-course analyses across SIV infection and ART were performed separately for total CD4⁺ T cells, CD4⁺ T cell subsets, and CD4⁻ T cells using the nonparametric Friedman test. Post hoc pairwise comparisons were performed using the Nemenyi test, with TSBH-FDR correction applied for multiple comparisons. Depending on the biological question, analyses compared post-infection or post-ART time points either directly to the pre-infection baseline (W0) or across all time points. Significantly altered metabolite–timepoint combinations (*q* ≤ 0.05) were overlaid on heatmaps as colored circles indicating the reference time point used in each comparison. Associations between MIST metabolic variables and virological measures or type I interferon transcriptional scores were assessed using Spearman rank correlation. For these analyses, only measurements matched within the same animal and the same time point were used to preserve biological pairing between metabolic and virological or transcriptional variables. Correlations used for chord diagram visualizations were calculated using Spearman rank correlation across samples to assess relationships between immune cell subset markers and metabolic marker gMFI values. Chord diagrams were generated using the circlize R package^24^ adapted from previously published MetFlowChord scripts, with link color representing the direction and magnitude of the Spearman correlation coefficient (−1 to +1) and link width proportional to the absolute correlation value. Statistical significance was denoted as **** for *q* ≤ 1×10⁻⁴, *** for *q* ≤ 1×10⁻³, ** for *q* ≤ 0.01, and * for *q* ≤ 0.05, unless otherwise indicated. For in vitro infection experiments, −*q* ≤ 0.10 was additionally indicated where specified. For correlations between MIST metabolic variables and virological measures, correlations meeting nominal significance (*p* ≤ 0.05) but not surviving FDR correction (*q* > 0.05) were indicated by a single dot (•), as specified in the corresponding figure legends. For experiments assessing the effects of ACC1 inhibition with Firsocostat, two-way repeated-measures ANOVA was used to test for the effects of activation status, treatment, and their interaction. Post hoc within-donor comparisons were performed using paired *t* tests, and corresponding Wilcoxon signed-rank test results were additionally reported in the supplementary data as a nonparametric reference supporting similar conclusions. Multiple testing across markers was corrected using the TSBH-FDR method.

## Data Availability Statement

All relevant data are included in the manuscript, supplementary and supplemental material. Values for all data points in graphs are reported in the Source Data file. All RNA sequencing data originating from this study have been deposited in NCBI GEO under the accession code: xx For reviewers: Enter token XX into the box.

## LIST OF SUPPLEMENTARY MATERIALS

Figures S1 to S13

Tables S1 to S5

Supplementary Data File 1

Source Data File

## General

We acknowledge the staff at the New Iberia Research Center for macaque samples processing, Leidos at NCI Frederick for plasma viral loads, the staff of the Flow Cytometry Core Facility at the Robert H. Lurie Comprehensive Cancer Center for assistance with cell sorting when needed and the staff of the Metabolomics Core of Northwestern University in Chicago.

## Authors contributions

JK and AA performed data analysis and contributed to writing the manuscript; KLY, RA, CAS, CAS, EBT contributed to sample processing and assays; MA, DB and FJV coordinated sample collection from the NHP study; IF and UN contributed to data interpretation and writing the manuscript; EM conceptualized the studies, analyzed the data, and wrote the manuscript.

## Declaration of generative AI

In the preparation of this manuscript the authors used ChatGPT 5.2 or Claude 3.7 Sonnet to improve readability of the text. After using this tool, the authors reviewed and edited the content, as necessary. The authors take full responsibility for the content of the publication.

## References

1. Kang, S. & Tang, H. HIV-1 Infection and Glucose Metabolism Reprogramming of T Cells: Another Approach Toward Functional Cure and Reservoir Eradication. Front Immunol 11, 572677 (2020).

2. Saez-Cirion, A. & Sereti, I. Immunometabolism and HIV-1 pathogenesis: food for thought. Nat Rev Immunol 21, 5–19 (2021).

3. Clerc, I. et al. Entry of glucose- and glutamine-derived carbons into the citric acid cycle supports early steps of HIV-1 infection in CD4 T cells. Nat Metab 1, 717–730 (2019).

4. Taylor, H.E. et al. mTOR Overcomes Multiple Metabolic Restrictions to Enable HIV-1 Reverse Transcription and Intracellular Transport. Cell Rep 31, 107810 (2020).

5. Valle-Casuso, J.C. et al. Cellular Metabolism Is a Major Determinant of HIV-1 Reservoir Seeding in CD4(+) T Cells and O[ers an Opportunity to Tackle Infection. Cell Metab 29, 611–626 e615 (2019).

6. Rasheed, S., Yan, J.S., Lau, A. & Chan, A.S. HIV replication enhances production of free fatty acids, low density lipoproteins and many key proteins involved in lipid metabolism: a proteomics study. PLoS One 3, e3003 (2008).

7. Abrahams, M.R. et al. The replication-competent HIV-1 latent reservoir is primarily established near the time of therapy initiation. Sci Transl Med 11 (2019).

8. Pankau, M.D. et al. Dynamics of HIV DNA reservoir seeding in a cohort of superinfected Kenyan women. PLoS Pathog 16, e1008286 (2020).

9. Shahid, A. et al. The replication-competent HIV reservoir is a genetically restricted, younger subset of the overall pool of HIV proviruses persisting during therapy, which is highly genetically stable over time. J Virol 98, e0165523 (2024).

10. Gantner, P. et al. HIV rapidly targets a diverse pool of CD4(+) T cells to establish productive and latent infections. Immunity 56, 653–668 e655 (2023).

11. Butler, K.M. et al. Rapid viral rebound after 4 years of suppressive therapy in a seronegative HIV-1 infected infant treated from birth. Pediatr Infect Dis J 34, e48–51 (2015).

12. Brooks, K. et al. HIV-1 variants are archived throughout infection and persist in the reservoir. PLoS Pathog 16, e1008378 (2020).

13. Brodin, J. et al. Establishment and stability of the latent HIV-1 DNA reservoir. Elife 5 (2016).

14. Brooks, K. et al. Proviral Turnover During Untreated HIV Infection Is Dynamic and Variable Between Hosts, Impacting Reservoir Composition on ART. Front Microbiol 12, 719153 (2021).

15. Munger, J. et al. Systems-level metabolic flux profiling identifies fatty acid synthesis as a target for antiviral therapy. Nat Biotechnol 26, 1179–1186 (2008).

16. Kulkarni, M.M. et al. Cellular fatty acid synthase is required for late stages of HIV-1 replication. Retrovirology 14, 45 (2017).

17. Aliyari, S.R. et al. Suppressing fatty acid synthase by type I interferon and chemical inhibitors as a broad spectrum anti-viral strategy against SARS-CoV-2. Acta Pharm Sin B 12, 1624–1635 (2022).

18. Kanno, T. et al. SCD2-mediated monounsaturated fatty acid metabolism regulates cGAS-STING-dependent type I IFN responses in CD4(+) T cells. Commun Biol 4, 820 (2021).

19. Berod, L. et al. De novo fatty acid synthesis controls the fate between regulatory T and T helper 17 cells. Nat Med 20, 1327–1333 (2014).

20. Endo, Y. et al. ACC1 determines memory potential of individual CD4(+) T cells by regulating de novo fatty acid biosynthesis. Nat Metab 1, 261–275 (2019).

21. Guo, H. et al. Multi-omics analyses reveal that HIV-1 alters CD4(+) T cell immunometabolism to fuel virus replication. Nat Immunol 22, 423–433 (2021).

22. Ambikan, A.T. et al. Genome-scale metabolic models for natural and long-term drug-induced viral control in HIV infection. Life Sci Alliance 5 (2022).

23. Ambikan, A.T. et al. Multi-omics personalized network analyses highlight progressive disruption of central metabolism associated with COVID-19 severity. Cell Syst 13, 665–681 e664 (2022).

24. Ahl, P.J. et al. Met-Flow, a strategy for single-cell metabolic analysis highlights dynamic changes in immune subpopulations. Commun Biol 3, 305 (2020).

25. Lee, J. et al. Regulator of fatty acid metabolism, acetyl coenzyme a carboxylase 1, controls T cell immunity. J Immunol 192, 3190–3199 (2014).

26. Patsoukis, N. et al. PD-1 alters T-cell metabolic reprogramming by inhibiting glycolysis and promoting lipolysis and fatty acid oxidation. Nat Commun 6, 6692 (2015).

27. Frauwirth, K.A. & Thompson, C.B. Regulation of T lymphocyte metabolism. J Immunol 172, 4661–4665 (2004).

28. Michalek, R.D. et al. Peroxiredoxin II regulates e[ector and secondary memory CD8+ T cell responses. J Virol 86, 13629–13641 (2012).

29. Salmond, R.J., Brownlie, R.J., Meyuhas, O. & Zamoyska, R. Mechanistic Target of Rapamycin Complex 1/S6 Kinase 1 Signals Influence T Cell Activation Independently of Ribosomal Protein S6 Phosphorylation. J Immunol 195, 4615–4622 (2015).

30. Ju, S.T. et al. Fas(CD95)/FasL interactions required for programmed cell death after T-cell activation. Nature 373, 444–448 (1995).

31. Richards, H., Longhi, M.P., Wright, K., Gallimore, A. & Ager, A. CD62L (L-selectin) down-regulation does not a[ect memory T cell distribution but failure to shed compromises anti-viral immunity. J Immunol 180, 198–206 (2008).

32. Wang, S. et al. An atlas of immune cell exhaustion in HIV-infected individuals revealed by single-cell transcriptomics. Emerg Microbes Infect 9, 2333–2347 (2020).

33. Pollara, J. et al. Single-cell analysis of immune cell transcriptome during HIV-1 infection and therapy. BMC Immunol 23, 48 (2022).

34. Kanno, T. et al. SCD2-mediated cooperative activation of IRF3-IRF9 regulatory circuit controls type I interferon transcriptome in CD4(+) T cells. Front Immunol 13, 904875 (2022).

35. Stuve, P. et al. ACC1 is a dual metabolic-epigenetic regulator of Treg stability and immune tolerance. Mol Metab 94, 102111 (2025).

36. Fleta-Soriano, E. et al. The Myxobacterial Metabolite Soraphen A Inhibits HIV-1 by Reducing Virus Production and Altering Virion Composition. Antimicrob Agents Chemother 61 (2017).

37. Vanhamel, J., Bruggemans, A. & Debyser, Z. Establishment of latent HIV-1 reservoirs: what do we really know? J Virus Erad 5, 3–9 (2019).

38. Okoye, A.A. et al. Early antiretroviral therapy limits SIV reservoir establishment to delay or prevent post-treatment viral rebound. Nat Med 24, 1430–1440 (2018).

39. Mdluli, T. et al. Acute HIV-1 infection viremia associate with rebound upon treatment interruption. Med 3, 622–635 e623 (2022).

40. Passaes, C. et al. Early antiretroviral therapy favors post-treatment SIV control associated with the expansion of enhanced memory CD8(+) T-cells. Nat Commun 15, 178 (2024).

41. Moller, S.H., Hsueh, P.C., Yu, Y.R., Zhang, L. & Ho, P.C. Metabolic programs tailor T cell immunity in viral infection, cancer, and aging. Cell Metab 34, 378–395 (2022).

42. Bowman, E.R. et al. In Vitro Exposure of Leukocytes to HIV Preexposure Prophylaxis Decreases Mitochondrial Function and Alters Gene Expression Profiles. Antimicrob Agents Chemother 65 (2020).

43. Angin, M. et al. Metabolic plasticity of HIV-specific CD8(+) T cells is associated with enhanced antiviral potential and natural control of HIV-1 infection. Nat Metab 1, 704–716 (2019).

44. Perdomo-Celis, F. et al. Impact of rosuvastatin on the memory potential and functionality of CD8(+) T cells from people with HIV. EBioMedicine 114, 105672 (2025).

45. Ahmed, D., Roy, D. & Cassol, E. Examining Relationships between Metabolism and Persistent Inflammation in HIV Patients on Antiretroviral Therapy. Mediators Inflamm 2018, 6238978 (2018).

46. Alzahrani, J. et al. Inflammatory and immunometabolic consequences of gut dysfunction in HIV: Parallels with IBD and implications for reservoir persistence and non-AIDS comorbidities. EBioMedicine 46, 522–531 (2019).

47. Rodriguez, N.R. et al. Oxidative phosphorylation in HIV-1 infection: impacts on cellular metabolism and immune function. Front Immunol 15, 1360342 (2024).

48. Steinert, E.M., Vasan, K. & Chandel, N.S. Mitochondrial Metabolism Regulation of T Cell-Mediated Immunity. Annu Rev Immunol 39, 395–416 (2021).

49. Klein, S.L. & Flanagan, K.L. Sex di[erences in immune responses. Nat Rev Immunol 16, 626–638 (2016).

50. Mauvais-Jarvis, F. Sex di[erences in energy metabolism: natural selection, mechanisms and consequences. Nat Rev Nephrol 20, 56–69 (2024).

51. Fray, E.J. et al. Antiretroviral therapy reveals triphasic decay of intact SIV genomes and persistence of ancestral variants. Cell Host Microbe 31, 356–372 e355 (2023).

52. Ewels, P.A. et al. The nf-core framework for community-curated bioinformatics pipelines. Nat Biotechnol 38, 276–278 (2020).

53. Di Tommaso, P. et al. Nextflow enables reproducible computational workflows. Nat Biotechnol 35, 316–319 (2017).

54. Gruning, B. et al. Bioconda: sustainable and comprehensive software distribution for the life sciences. Nat Methods 15, 475–476 (2018).

55. da Veiga Leprevost, F., et al. BioContainers: an open-source and community-driven framework for software standardization. Bioinformatics 33, 2580–2582 (2017).

56. Dobin, A., et al. STAR: ultrafast universal RNA-seq aligner. Bioinformatics 29, 15–21 (2013).

57. Patro, R., Duggal, G., Love, M.I., Irizarry, R.A. & Kingsford, C. Salmon provides fast and bias-aware quantification of transcript expression. Nat Methods 14, 417–419 (2017).

58. Fang, Z., Liu, X. & Peltz, G. GSEApy: a comprehensive package for performing gene set enrichment analysis in Python. Bioinformatics 39 (2023).

59. Kuleshov, M.V. et al. Enrichr: a comprehensive gene set enrichment analysis web server 2016 update. Nucleic Acids Res 44, W90–97 (2016).

60. Foroutan, M., et al. Single sample scoring of molecular phenotypes. BMC Bioinformatics 19, 404 (2018).

61. Wang, H., et al. Genome-scale metabolic network reconstruction of model animals as a platform for translational research. Proc Natl Acad Sci U S A 118 (2021).

62. Robinson, J.L., et al. An atlas of human metabolism. Sci Signal 13 (2020).

63. Wang, H., et al. RAVEN 2.0: A versatile toolbox for metabolic network reconstruction and a case study on Streptomyces coelicolor. PLoS Comput Biol 14, e1006541 (2018).

64. Gustafsson, J., et al. Generation and analysis of context-specific genome-scale metabolic models derived from single-cell RNA-Seq data. Proc Natl Acad Sci U S A 120, e2217868120 (2023).

65. Zheng, G.X., et al. Massively parallel digital transcriptional profiling of single cells. Nat Commun 8, 14049 (2017).

66. Hao, Y., et al. Dictionary learning for integrative, multimodal and scalable single-cell analysis. Nat Biotechnol 42, 293-304 (2024).

