## Supplementary Figures for "Type I IFN reprograms CD4⁺ T cell lipid metabolism as an antiviral effector mechanism during acute HIV/SIV infection"

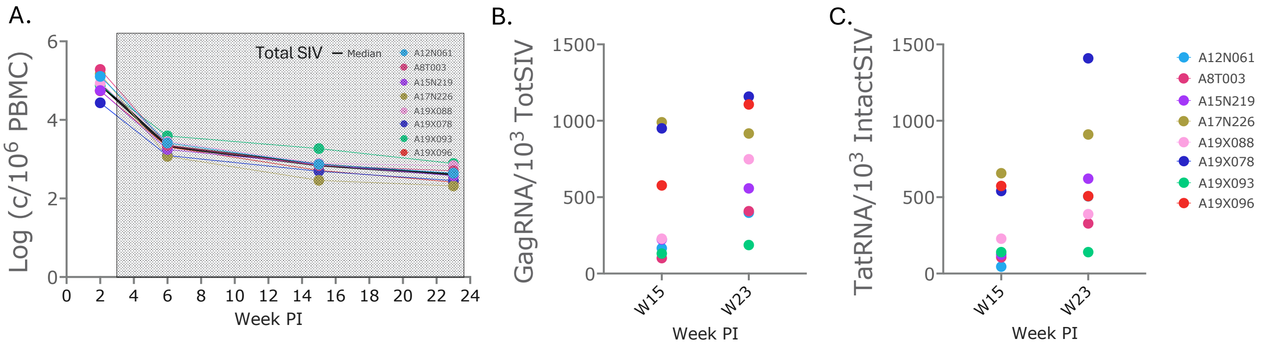


**Figure S1. Cell associated viral DNA and RNA in PBMC at specified time points post infection.** A) Cell associated total SIV copies per million PBMC B) Cell associated Gal RNA levels (gag copies) normalized for total DNA levels C) Cell associated copies of Tat/Rev multiply spliced RNA normalized for copies of intact provirus.

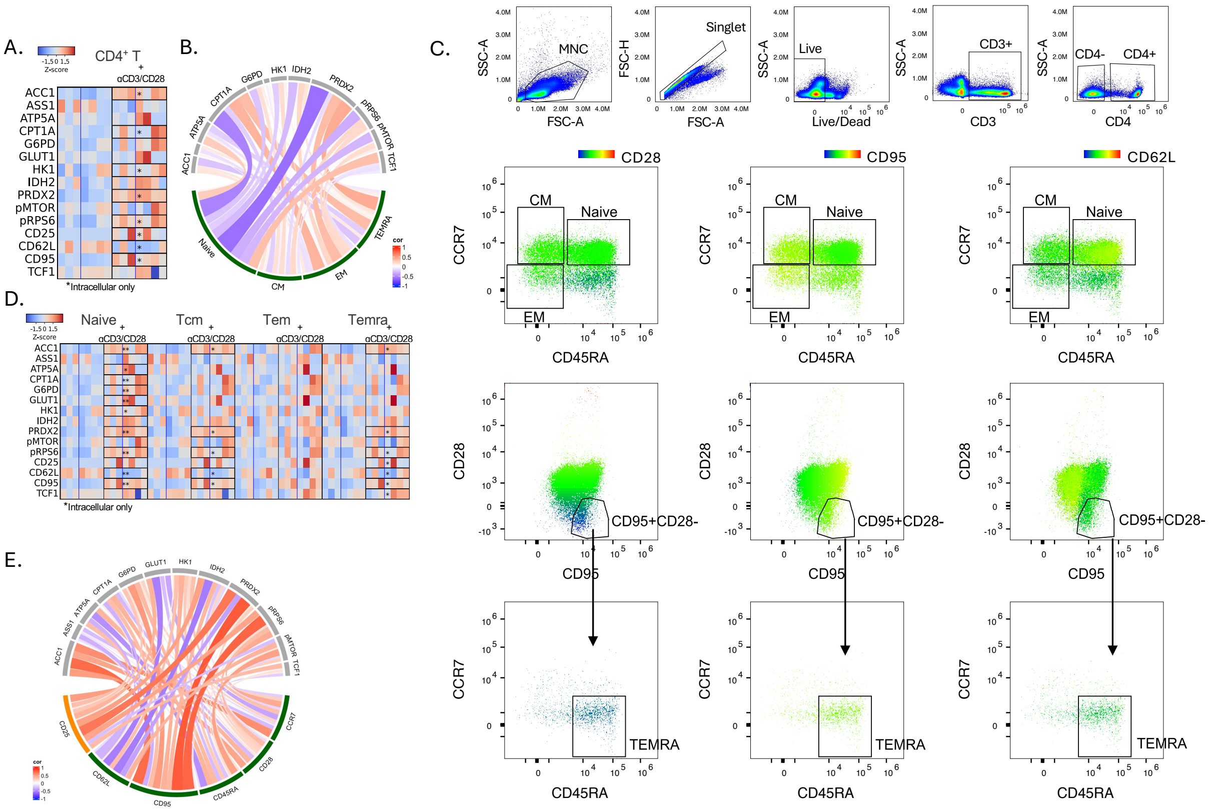


**Figure S2. In vitro MIST Validation.** A) Heatmap of levels (MFI) of each MIST metabolic variable within CD4$^+$ T cells of rhesus isolated CD4 T cells (n=3) and PBMC (n=4) are shown comparing activated (anti-CD3/CD28) to non-activated cells cultured in parallel (*q≤0.05, **q≤0.01, ***q≤1e-3 , and ****q≤1e-4 by Wilcoxon signed-rank test with TSBH-FDR correction). B) Chord diagram showing Spearman correlations between CD4$^+$ T cell subsets and metabolic marker gMFI levels. Links represent the strength of association between subtype membership and each metabolic variable, with color indicating the direction and magnitude of the correlation (–1 to +1) and link width proportional to the absolute correlation value. C) Representative flow cytometry plots showing the gating strategy used to define CD4$^+$ T cell subsets. Lymphocytes were first identified based on forward scatter (FSC) versus side scatter (SSC). Single cells were gated using FSC-A versus FSC-H, followed by exclusion of dead cells based on FVS440UV negative staining. Live singlet cells were then gated on $CD3^+CD4^+$ cells to define total $CD4^+$ T cells. Within the $CD4^+$ T cell population, subsets were defined based on CD45RA and CCR7 expression as follows: naïve T cells ($CD45RA^+$ $CCR7^+$), central memory (CM; $CD45RA^-$ $CCR7^+$), and effector memory (EM; $CD45RA^-$ $CCR7^-$). TEMRA cells were defined within the $CD4^+$ T cell population as $CD95^+CD28^-$ cells and further gated as $CD45RA^+$ $CCR7^-$. The three columns represent the same representative sample, with fluorescence intensity displayed for different markers: CD28 (left), CD95 (middle), and CD62L (right). Color scale indicates marker expression intensity. D) Comparison of levels of MIST metabolic variables between activated and non-activated cells within each CD4 T cell subset. (*q≤0.05, **q≤0.01 by Wilcoxon signed-rank test with TSBH-FDR correction). E) Chord diagram showing Spearman correlations between MIST variables levels in macaque PBMC (fresh/non activated) and activation markers. Links represent the strength of association, while color indicates direction and magnitude of the correlation (–1 to +1).

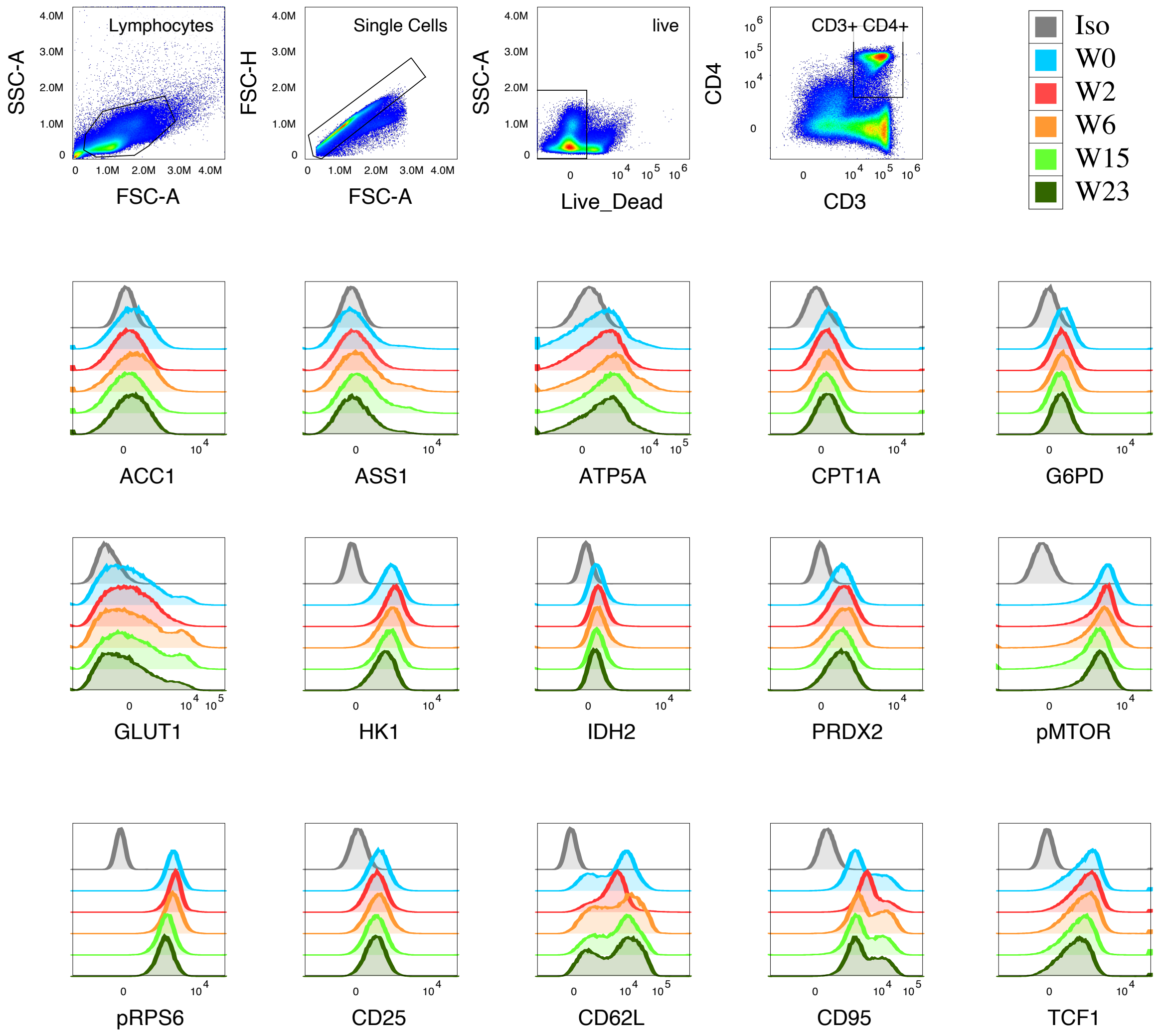


**Figure S3. Representative hating strategy and histogram overlays for MIST profiling of CD4⁺ T cells during SIV infection and ART across study time points.**

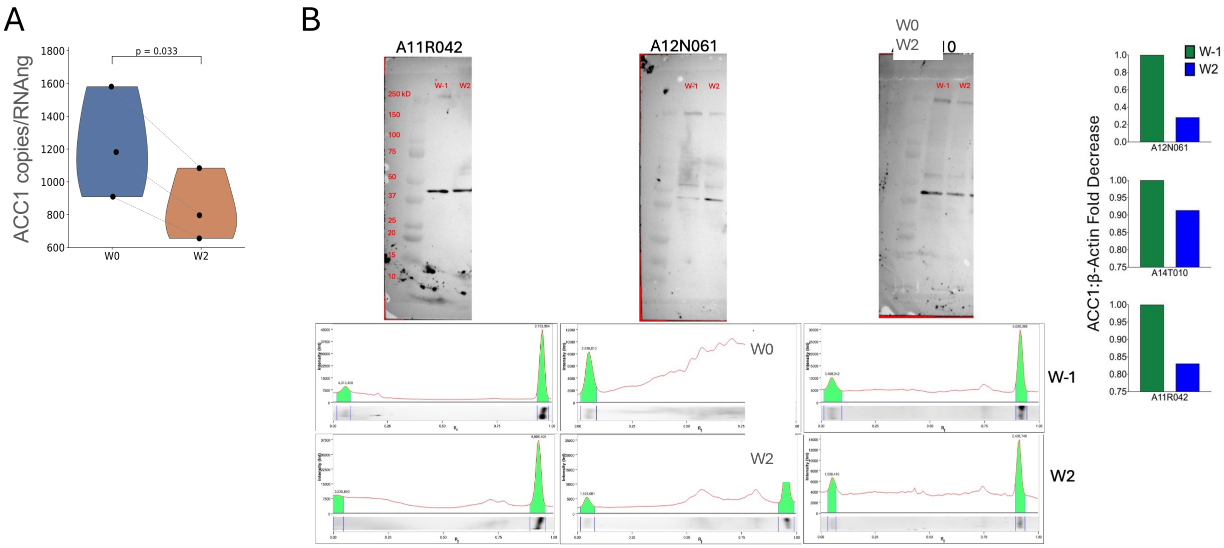


**Figure S4. SIV infection decreases ACC1 in CD4+ T cells at peak infection. A)** Copies of ACC1 mRNA per ng of CD4+ T cells RNA left over from 3 of the 6 macaques used for RNAseq analysis. Copies of ACC1 were analyzed by dPCR (Absolute Q by Thermo Fisher) and normalized on amount of RNA (measured by Qbit) used for the RT reaction. (Paired t test; $\alpha$=p<0.05) **B)** Densitometric quantification analysis (below) of western blots (above) from three macaques showing ACC1 and $\beta$-actin expression levels. CD4+ T cells were sorted from PBMC taken pre-infection (W0) and two weeks post-infection (W2). An equal number of sorted cells from each time point were lysed and ran on a 4-15% density gradient SDS-PAGE gel before transfer to a nitrocellulose membrane. Membrane was incubated with primary ACC1 and $\beta$-actin antibodies overnight, and with HRP-conjugated secondary antibody for one hour. The membrane was then exposed to SuperSignal WestFemto HRP substrate and visualized using an iBright chemiluminescent imaging system. Signal was quantified using densiometric analysis in BioRad ImageLab software and used to determine the ratio of ACC1 to $\beta$-actin signal intensity for each time point. ACC1:$\beta$-actin intensity ratio from W0 was used as a baseline to determine fold decrease in W2 (right).

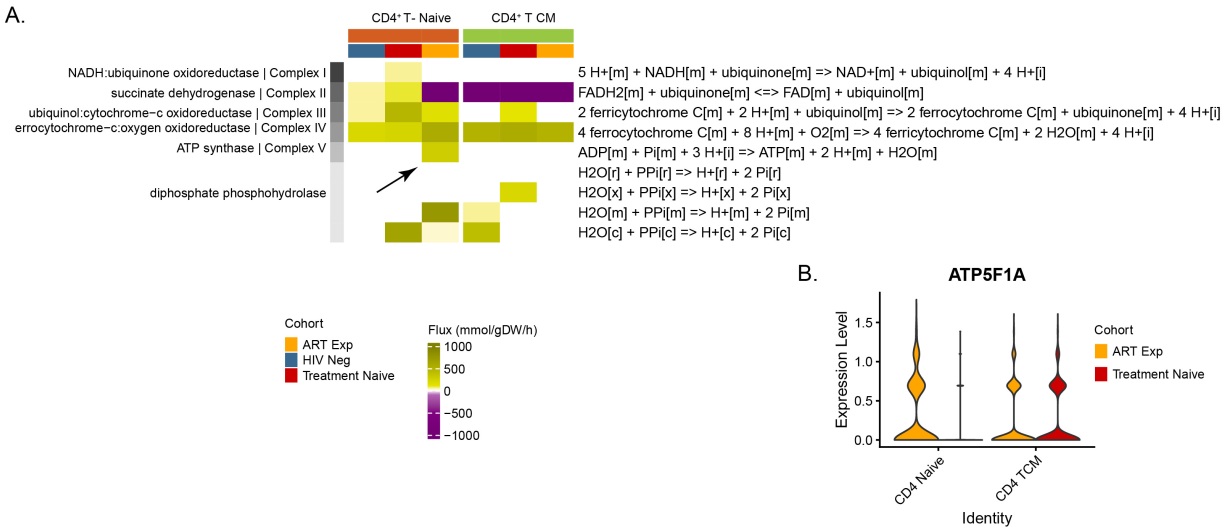


**Figure S5. OXPHOS is upregulated in CD4 T cells from PWH on ART compared to untreated and PWoH.** A) Heatmap showing flux values predicted by flux balance analysis for reactions involved in the oxidative phosphorylation pathway in CD4$^{+}$ naive and central memory T cells. Column annotations indicate cohort information, while row annotations denote the reaction names and reaction types. B) Violin plot illustrating the expression levels of ATP5F1A in CD4$^{+}$ naive and central memory T cells from ART-treated and untreated samples.

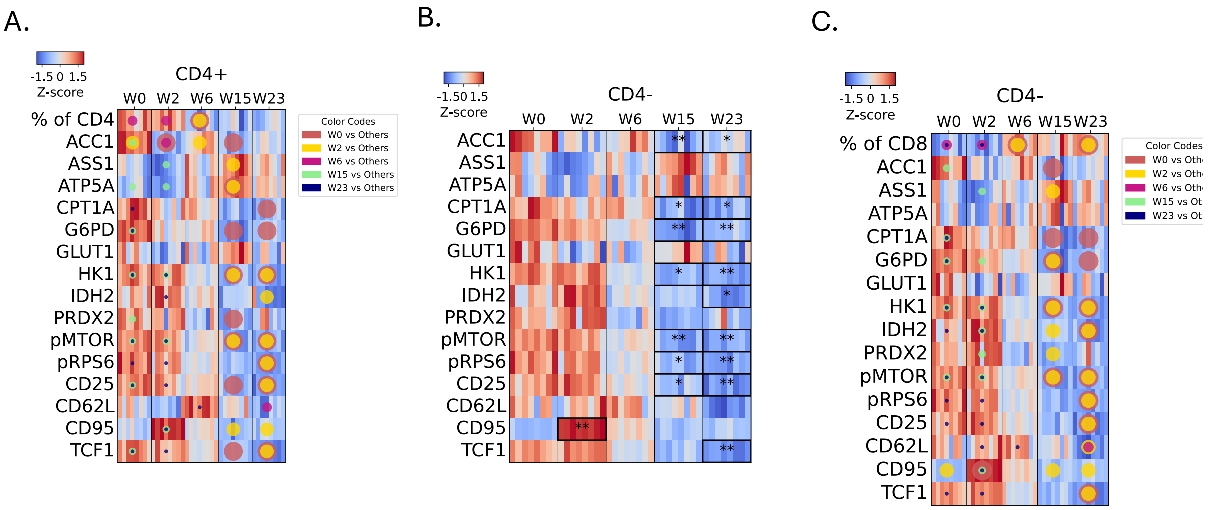


**Figure S6 Comparisons of MIST variables in macaque $CD4^+$ and $CD4^-$ T cells between all-time points.** A) Heatmap of MIST variables' levels in $CD4^+$ T cell at the indicated weeks, analyzed using the Friedman test with Nemenyi post-hoc and TSBH-FDR correction comparing each time point to all others. Significantly altered metabolite–timepoint combinations (q ≤ 0.05) are overlaid as colored circles, where circle color and size denote the baseline time point (W0, W2, W6, W15, W23) used in each pairwise comparison. B-C) Heatmaps of MIST variables levels within $CD4^-$ T cells are shown at time points post infection compared to before infection (B) With Friedman Test with Nemenyi posthoc and TSBH-FDR correction comparing each time point to before infection (*q≤0.05, **q≤0.01, ***q≤1e-3) (C) With Friedman test with Nemenyi post-hoc and TSBH-FDR correction comparing each time point to all others. Significantly altered metabolite–timepoint combinations (q ≤0.05) are overlaid as colored circles, where circle color and size denote the baseline time point (W0, W2, W6, W15, W23) used in each pairwise comparison.

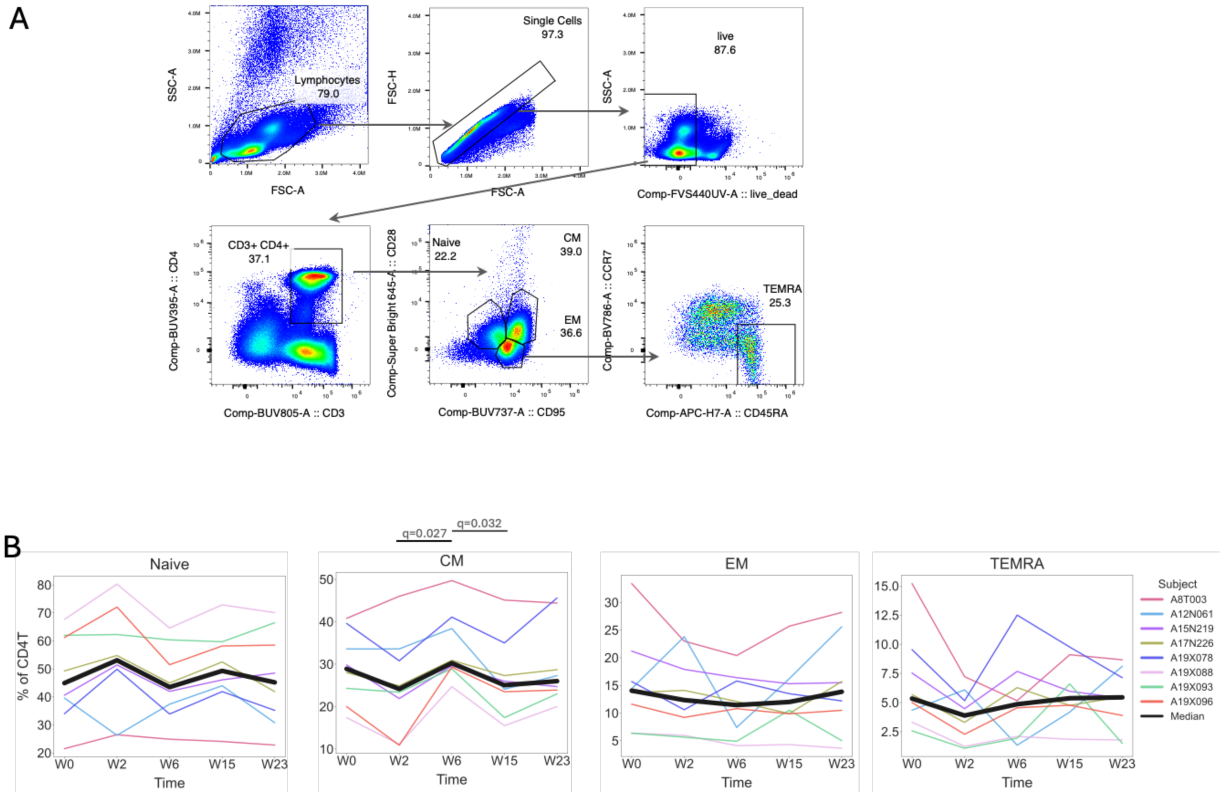


**Figure S7. CD4$^{+}$ T-cell subset gating and frequencies during SIV infection and ART.** A) Representative gating strategy used to identify CD3$^{+}$ CD4$^{+}$ T cells and define naïve, central memory (CM), effector memory (EM), and TEMRA subsets for MIST analysis of ex vivo PBMC. B) Frequencies of CD4$^{+}$ T-cell subsets across the different time points in the MIST assay were compared by Friedman test with Nemenyi post-hoc and TSBH-FDR correction comparing each time point to all others within a given subset. Significant comparisons (q≤0.05) are shown.

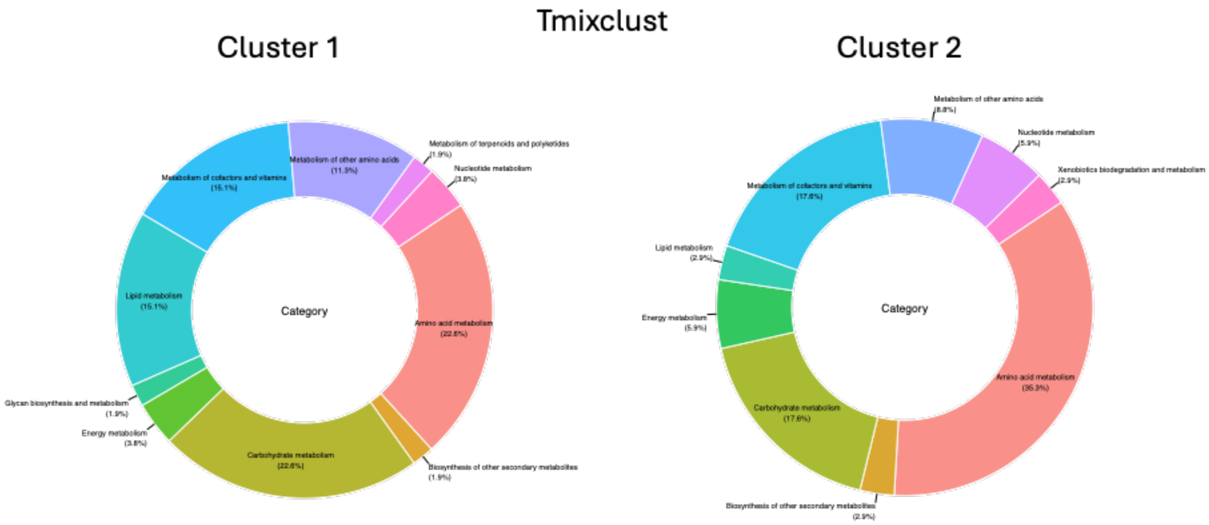


**Figure S8 Proportion of metabolites categories in each cluster after Tmixclust analysis of plasma metabolomics at weeks 0, 2, 6 and 23 post infection.**

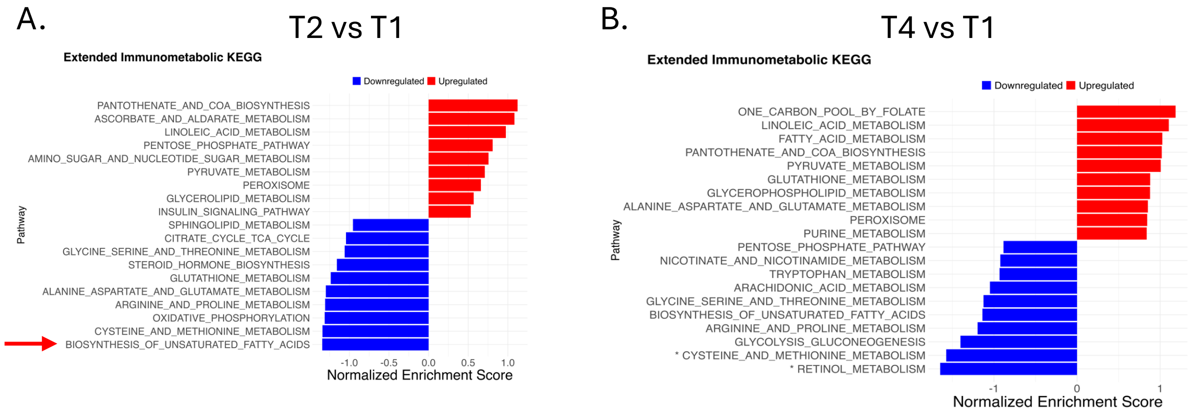


**Figure S9. GSEA performed on genes ranked by $\log_2$FC from the T2 vs T1 and T4 vs T1 comparisons using the Immunometabolic KEGG gene sets.**

GSEA performed on RNAseq data from sorted CD4+ T cells at indicated time points were analyzed for enrichment of metabolic pathways included in the immunometabolic KEGG gene set (ED Table S3) (Adj p≤0.05).

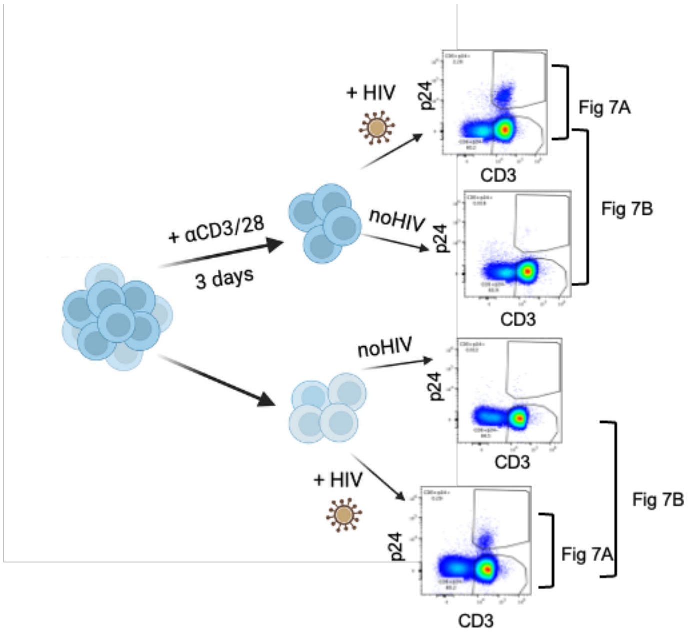


**Figure S10. Schematic of in vitro HIV infection experiment and population compared in Fig 7 with gating strategy.**

A.

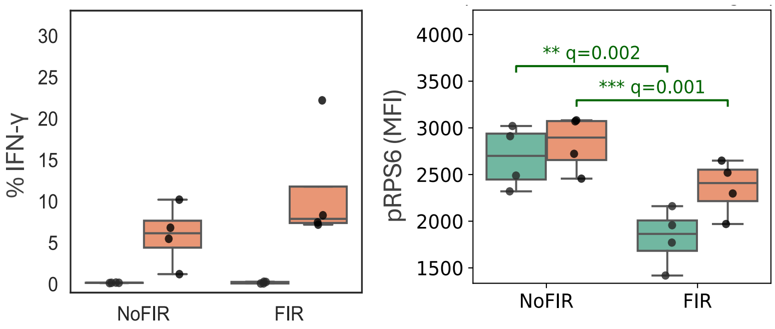


B.

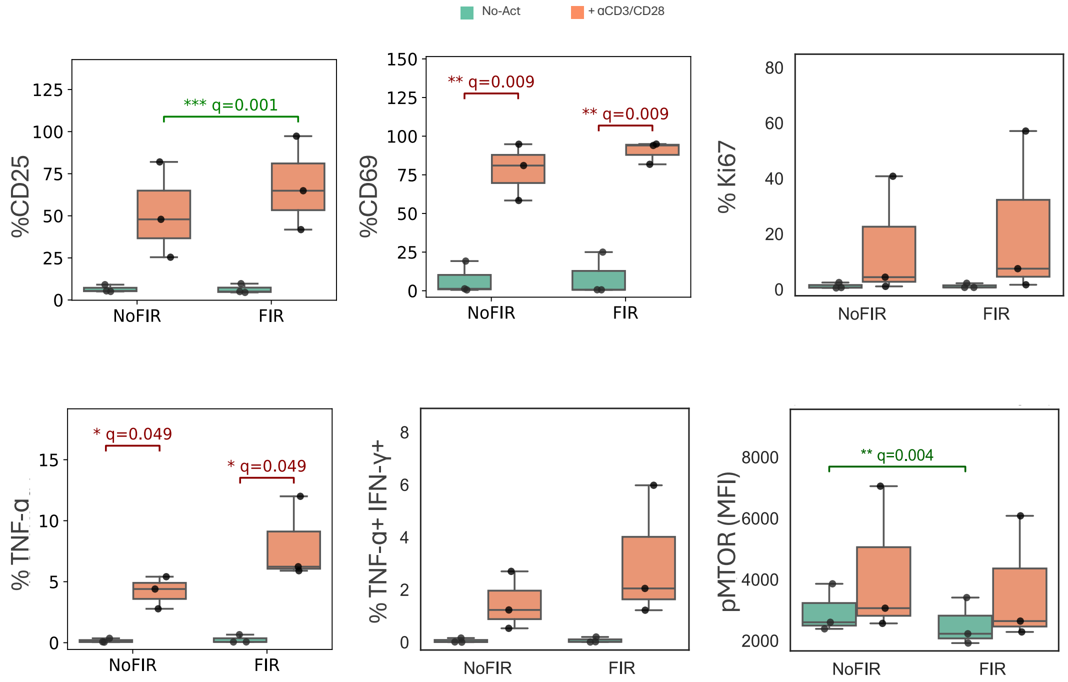


**Figure S11 Blocking ACC1 with Firsocostat (Fir) increases response to CD3/28 activation in rhesus and human CD4 T cells.** A-B) Rhesus PBMC (A) and Human PBMC (B) were activated on plate coated with anti-CD3 and anti-CD28 for 48hrs with 100U/ml of IL2 or left in culture in parallel in absence of activation with 20U/ml of IL2. The increase in frequency of IFN-g and phspho-(Ser235/236) RPS6 are shown for rhesus cells (A). The increase in frequency of $CD25^+$, $CD69^+$ and $Ki67^+$ human $CD4^+$ T cells and in cells producing TNF-α, TNF-α and IFN-γ and the levels of phospho- (activated)MTOR in activated (orange) compared to non-activated (green) cells in presence and absence of Fir is shown. (Two-way repeated-measures ANOVA with paired post hoc t-tests was used, and multiple testing was corrected using the FDR-TSBH method;

*q≤0.05, **q≤0.01, ***q≤1e-3. Similar conclusions were reached with non-parametric tests. See Supplementary Data File)

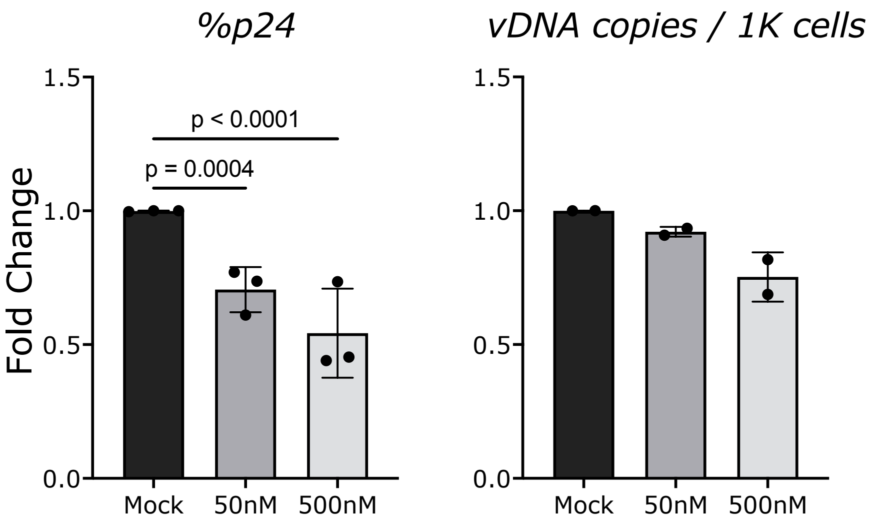


**Figure S12 Blocking ACC1 with Firsocostat (Fir) decreases HIV infection in vitro in CD4$^+$ T cells.** Isolated CD4$^+$ T cells from 3 donors were infected by spinoculation and cultured in presence or absence of the indicated concentration of Fir in triplicates or quadruplicates per donor and condition. A) HIV-p24 levels were measured intracellularly after 4-7 days of infection and compared by mixed-effect analysis (including all replicates) and BH-FDR p is shown. Shown are the means of the replicates for each donor and the bar represents the SD of the means. B) For confirmation, we include the cell-associated vDNA (HIV-gag) levels per 1000 cells from 2 of the 3 donors. Shown are the means of the replicates for each donor and the bar represents the SD of the means.

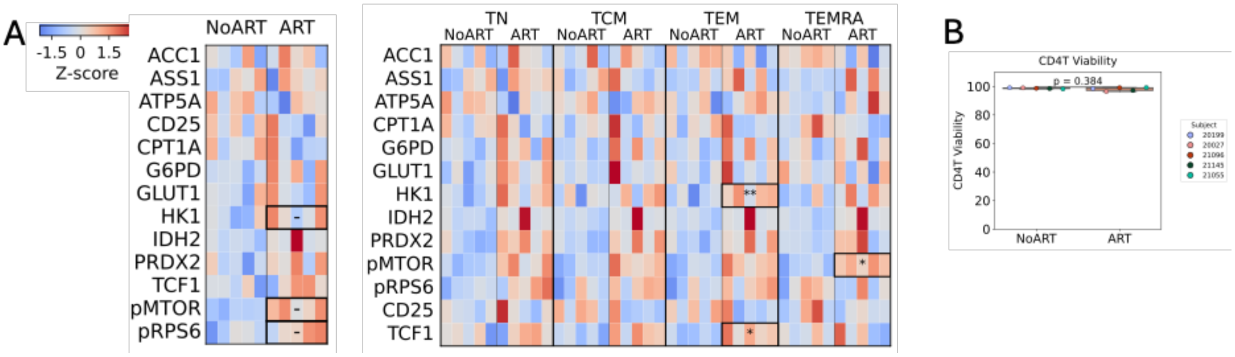


**Figure S13. In vitro treatment of macaque PBMC with ART drugs does not recapitulate metabolic changes in vivo.** PBMC from 5 different macaques were thawed and cultured for 7 days in RPMI 10% FBS 10U/ml of IL2 and ART (TDF 1μM, FTC 5μM and DGT 5μM expected peak plasma concentrations of in vivo administered drugs) was added to half of the cultures every 2-3 days. On day 8, the level of metabolic variables was assessed by MIST staining. A) No significant differences (q<0.05) were detected by direct comparison of the MIST variables in gated live, CD3$^+$ CD4$^+$T cells in ART-treated compared to mock treated cells. Significant changes in HK and TCF1 levels were noted in gated TEM cells (CD95$^+$ CD28$^-$) and PRDX2 in TEMRA (CD95$^+$ CD28$^-$ /CD45RA$^+$ CCR7$^-$). Paired t-test FDR corrected - q<0.1; significance q*<0.01; **<0.01. B) No changed in cell viability were noted by live/dead exclusion staining (Paired t-test, two-sided $\alpha$=0.05).

### Supplementary Tables

**Table S1 MIST1 Panel List of Markers and Antibodies**

**Table S2 Basic KEGG Metabolic Pathways.** This table lists all the genes in each KEGG pathway included in this collection. This collection includes only major metabolic pathways known to have a major role in immune cells.

**Table S3 Extended KEGG Metabolic Pathways** This table lists all the genes in each KEGG pathway included in this collection. This collection includes all metabolic pathways known to have a role in immune cells.

**Table S4 MIST2 Panel List of Markers and Antibodies**

**Table S5. List of IFN-stimulated genes used to determine the IFN-I scores.**

**Supplementary Data File. Parametric and non-parametric tests results for Figure 7 A, B, E**
